# Srs2 helicase prevents the formation of toxic DNA damage during late prophase I of yeast meiosis

**DOI:** 10.1101/518035

**Authors:** Hiroyuki Sasanuma, Hana Subhan M. Sakurai, Yuko Furihata, Kiran Challa, Lira Palmer, Susan M. Gasser, Miki Shinohara, Akira Shinohara

## Abstract

Proper repair of double-strand breaks (DSBs) is key to ensure proper chromosome segregation. In this study, we found that the deletion of the *SRS2* gene, which encodes a DNA helicase necessary for the control of homologous recombination, induces aberrant chromosome segregation during budding yeast meiosis. This abnormal chromosome segregation in *srs2* cells accompanies the formation of a novel DNA damage induced during late meiotic prophase-I. The damage may contain long stretches of single-stranded DNAs (ssDNAs), which lead to aggregate formation of a ssDNA binding protein, RPA, and a RecA homolog, Rad51, as well as other recombination proteins inside of the nuclei. The Rad51 aggregate formation in the *srs2* mutant depends on the initiation of meiotic recombination and occurs in the absence of chromosome segregation. Importantly, as an early recombination intermediate, we detected a thin bridge of Rad51 between two Rad51 foci or among the foci in the *srs2* mutant, which is rarely seen in wild type. These might be cytological manifestation of the connection of two DSB ends and multi-invasion. The DNA damage with Rad51 aggregates in the *srs2* mutant is passed through anaphase-I and -II, suggesting the absence of DNA damage-induced cell-cycle arrest after the pachytene stage. We propose that Srs2 helicase resolves early protein-DNA recombination intermediates to suppress the formation of aberrant lethal DNA damage during late prophase-I.

## Introduction

In sexually reproducing organisms, meiosis, a specialized form of cell division, produces haploid gametes from diploid germ cells. Following DNA replication, reciprocal recombination takes place to connect the homologous chromosomes and to generate genetic diversity of gametes. With arm cohesion, the connection between the chromosomes, which is cytologically visualized as chiasma, is essential for faithful chromosome segregation during meiosis I by antagonizing the pulling force by spindle microtubules to create tension (Petronczki et al. 2003).

Meiotic recombination is initiated by the generation of DNA double-strand breaks (DSBs) by a meiosis-specific topoisomerase-like protein, Spo11, at recombination hotspots (Keeney et al. 1997). Subsequently, the end of DSBs is quickly resected to produce 3’-overhanging single-stranded DNAs (ssDNAs). Replication protein A (RPA) binds to the ssDNAs, followed by the loading of Rad51, a homolog of bacterial RecA (Shinohara et al. 1992), with the assistance of auxiliary proteins, such as Rad52, Rad55-Rad57 and Pys3-Csm2-Shu1-Shu2 (a.k.a. Shu) (New et al. 1998; Sasanuma et al. 2013b; Shinohara and Ogawa 1998; Sung 1997). Rad51 filaments on ssDNA are active protein machinery for DNA homology search and strand exchange (Ogawa et al. 1993; Sung 1994). Rad51 filament activity is helped by Rad54, which belongs to the SNF2/SWI2 DNA helicase family (Shinohara et al. 1997b).

Whereas Rad51 is sufficient for the homolog search in recombination during mitosis, meiosis requires a meiosis-specific RecA homolog, Dmc1, for the recombination (Bishop et al. 1992). Dmc1 is essential for homology search/strand exchange in inter-homolog recombination during meiosis while Rad51 plays an auxiliary role by assisting Dmc1 assembly (Bishop 1994; Cloud et al. 2012; Shinohara et al. 1997a). Indeed, the Rad51 activity for inter-sister recombination during meiosis is suppressed by the action of a meiosis-specific Rad51 inhibitor, Hed1 (Tsubouchi and Roeder 2004). Like Rad51, Dmc1 forms a nucleo-protein filament on ssDNAs to catalyze the strand invasion of the DNA into its homologous duplex DNA for the formation of an intermediate, D(displacement)-loop (Hong et al. 2001).

In D-loop, DNA synthesis occurs from 3’-end of invading strand as a primer. When the synthesized DNA strand is ejected from the D-loop (Allers and Lichten 2001; Hunter and Kleckner 2001), the ejected synthesized ssDNA is able to anneal with the complementary ssDNA in the other end of the DSB. Annealing induces the second DNA synthesis to complete the recombination by producing non-crossovers. This pathway is called synthesis-dependent strand annealing (SDSA) (Allers and Lichten 2001). On the other hand, when the newly synthesized DNA is stably bound to the D-loop, ongoing DNA synthesis can extend a D-loop with a large displaced ssDNA, which is able to anneal with ssDNA on the opposite DSB ends. Additional processing of the intermediates leads to the formation of double-Holliday junction (dHJ) (Schwacha and Kleckner 1994). dHJs are specifically resolved into crossovers. Importantly, meiotic recombination is tightly coupled with chromosome morphogenesis such as the formation of the synaptonemal complex (SC), a meiosis-specific zipper-like chromosome structure, which juxtaposes homologous chromosomes in near vicinity (Cahoon and Hawley 2016).

Srs2 is a 3’-to-5’ SF1 helicase related to bacterial UvrD helicase (Rong et al. 1991). Srs2 protein has some distinct functional domains: 3’-5’ DNA helicase domain, Rad51-interaction domain, and also SUMO- and PCNA-binding domains in the C-terminus (Marini and Krejci 2010). Genetic analyses showed positive and negative roles of Srs2 in the recombination (Marini and Krejci 2010). Biochemical studies have demonstrated that purified Srs2 protein can dislodge Rad51 filament on ssDNAs and dramatically inhibits Rad51-joint molecules via direct interaction with Rad51 *in vitro* (Krejci et al. 2003; Veaute et al. 2003). This biochemical activity of Srs2 supports the idea of Srs2 function as an anti-recombinase. The Rad51-dismantling activity of Srs2 is confirmed by *in vivo* analysis (Sasanuma et al. 2013a).

Deletion of *SRS2* gene shows different kinds of genetic interaction with mutants deficient in DNA transaction. The *srs2Δ* is synthetic lethal with a mutation of the *SGS1*, encoding a RecQ-type DNA helicase. By forming a complex with Top3 and Rmi1, Sgs1 is known to dissolve the dHJ structure into noncrossovers (Cejka et al. 2010; Wu and Hickson 2003). Moreover, the *srs2Δ* is synthetic lethal with the deletion of the *RAD54* (Klein 2001), suggesting the role of Srs2 in a late stage of the recombination such as the post-invasion step in the recombination. This lethality is thought to be caused by a fatal defect in the resolution of toxic intermediates in the recombination process. This is supported by the fact that the deletion of *RAD51* can suppress the lethality of *srs2Δ sgs1Δ* and *srs2Δ rad54Δ* mutants (Gangloff et al. 2000; Schild 1995).

During mitosis, crossovers should be suppressed when DNA damage is spontaneously introduced, because the crossover between homologous chromosomes and sister chromatids results in the loss of heterozygosity. In contrast, as described above, meiotic recombination must give rise to at least one essential crossover per chromosome, which is fostered by a group of proteins called ZMM (Zip-, Msh-, Mer) (Shinohara et al. 2008). Previous genetic studies showed a role of Srs2 in meiosis (Palladino and Klein 1992; Sasanuma et al. 2013b). However, the molecular defects associated with *srs2* deletion in meiosis have not been described in detail. Therefore, it remains elusive how Srs2 regulates meiotic recombination.

In this study, we analyzed the role of Srs2 helicase in meiotic recombination, particularly looking at dynamics of its interacting partner, Rad51. We found that, in the absence of Srs2, abnormal DNA damage associated with Rad51 aggregation accumulates during late prophase-I, after the completion of meiotic recombination. The formation of this DNA damage in the *srs2* requires meiotic DSB formation, but is independent of chromosome segregation. We also detected thin line-staining of Rad51 connecting between two adjacent Rad51 foci in early prophase in the absence of Srs2, which is rarely seen in the wild type. We propose that Srs2 protects chromosomes in late meiotic prophase-I from accumulation of abnormal DNA damage by properly coupling the completion of meiotic recombination with chromosome morphogenesis.

## Materials and Methods

### Yeast strains and medium conditions

All yeast strains used in this article are isogenic derivatives of SK1 and listed in Table S1. *pCLB2-SGS1* and *RAD54-RFB* strains were a gift by Dr. Neil Hunter and Dr. Andreas Hochwagen, respectively. Mediums and culture conditions regarding meiosis are described in (Sasanuma et al. 2008).

### Antibodies and chemicals

The primary anti-sera were used as following concentrations; anti-Rad51 (guinea pig, 1/500), anti-Dmc1 (rabbit, 1/500), anti-Rad52 (rabbit, 1/300), anti-Rfa2 (rabbit, 1/500), anti-Zip1 (rabbit, 1/500), anti-Red1 (rabbit, 1/500), Anit-Hed1 (rabbit, 1/200) and anti-Mei5 (rabbit, 1/500) for cytology. Anti-Hed1 serum from rabbit was prepared for denatured Hed1 protein purified from *E. coli*. Anit-Nop1(mouse) is from Encor Biotech (MCA28-F2). α-tubulin is monoclonal antibody of rat that can recognize alpha subunit (AbD Serotec/BioRad, MCA77G). The second antibodies for staining were Alexa-fluor 488 (Goat) and 594 (Goat) IgG used at a 1/2000 dilution (Molecular Probes).

Rapamycin (LC-Laboratories, R-5000) and benomyl (methyl 1-[butylcarbamoyl]-2-benzimidazolecarbamate; Sigma Aldrich, PCode 1002355429) were dissolved in DMSO at a concentration of 1 mM and 30 mg/ml, respectively.

### Immuno-staining

Chromosome spreads were prepared using the Lipsol method as described previously (Shinohara et al. 2000; Shinohara et al. 2003). Immnostaining was conducted as described (Shinohara et al. 2000). Stained samples were observed using an epi-fluorescence microscope (BX51; Olympus, Japan) with a 100X objective (NA1.3). Images were captured by CCD camera (CoolSNAP; Roper, USA), and afterwards processed using IP lab and/or iVision (Sillicon, USA), and Photoshop (Adobe, USA) software tools.

### SIM imaging

The structured illumination microscopy was carried out using super resolution-structured illumination (SR-SIM) microscope (Elyra S.1 [Zeiss], Plan-Apochromat 63x/1.4 NA objective lens, EM-CCD camera [iXon 885; Andor Technology], and ZEN Blue 2010D software [Zeiss]) at Friedrich Miescher Institute for Biomedical Research, Switzerland. Image processing was performed with Zen software (Zeiss, Germany), NIH image J and Photoshop.

### Whole cell staining

Cells were fixed with 1/10 volume of 37% formaldehyde (Wako) and treated with 10 μg/ml Zymolyase 100T (Seikagaku) for 1.5 h. Cells were placed to the poly L-lysine (Sigma) coated slides and then fixed with cold 100% methanol, cold 100% acetone and cold 1X PBS. Slides were used for immuno-staning.

### Western Blotting

Western blotting was performed for cell lysates extracted by TCA method. After being harvested and washed twice with 20% TCA, cells were roughly disrupted by Yasui Kikai (Yasui Kikai Co Ltd, Japan). Protein precipitation recovered by centrifuge at 1600 *g* for 5min was suspended in SDS-PAGE sample buffer adjusting to pH8.8 and then boiled for 95°C, 2min.

### Southern Blotting

Southern blotting analysis was performed with the same procedure as in (Storlazzi et al. 1995). Genomic DNA prepared was digested with both *MluI* and *XhoI* (for crossover/non-crossover, upper panels) and *PstI* (for meiotic DSB, lower panels). Probes for Southern blotting were Probe “155” for crossover/non-crossover and Probe 291 for DSB detection as described in (Storlazzi et al. 1995). Image gauge software (Fujifilm Co. Ltd., Japan) was used for quantification for bands of R1, R3 and DSB I.

### Pulsed-field gel electrophoresis

For pulsed-field gel electrophoresis (PFGE), chromosomal DNA was prepared in agarose plugs as described in (Bani Ismail et al. 2014) and run at 14 °C in a CHEF DR-III apparatus (BioRad) using the field 6V/cm at a 120° angle. Switching times followed a ramp from 15.1 to 25.1 seconds. Durations of electrophoresis were 41 h for chromosome III.

### Statistics

Means ± S.D values are shown. Graphs were prepared using and Microsoft Excel and GraphPad Prism 7. Datasets were compared using the Mann-Whitney U-test. χ^2^-test was used for proportion.

## Results

### *SRS2* deletion markedly decreased spore viability

As reported previously (Palladino and Klein 1992; Sasanuma et al. 2013a), the *srs2* deletion mutant exhibits reduced spore viability of 36.8%, indicating a critical role of this helicase for meiosis (Fig. S1A). This marked reduction of the spore viability is somehow unexpected given a negative role of this helicase in recombination.

We also confirmed the kinetics of meiotic progression in *srs2Δ* strains by DAPI staining. In the wild-type strain, meiosis I started at 5 h after incubation with sporulation medium (SPM) and was sequentially followed by meiosis II. Finally, ~90% of the wild-type cells completed MII at around 8 h (Fig. 1A). In the *srs2Δ* mutant, the appearance of cells undergoing MI was delayed by ~2 h and ~75% of cells finished MII at 14 h (Fig. 1A). A similar delay was observed for a *srs2* mutant in a different strain background previously (Palladino and Klein 1992). This indicates a defect during prophase-I in the *srs2* mutant. In *srs2* cells after sporulation; e.g. 12 h, we often detected fragmented DAPI bodies in a cell/spore (Fig. 1B), indicating a defect in chromosome segregation during the mutant meiosis.

**Figure 1.**
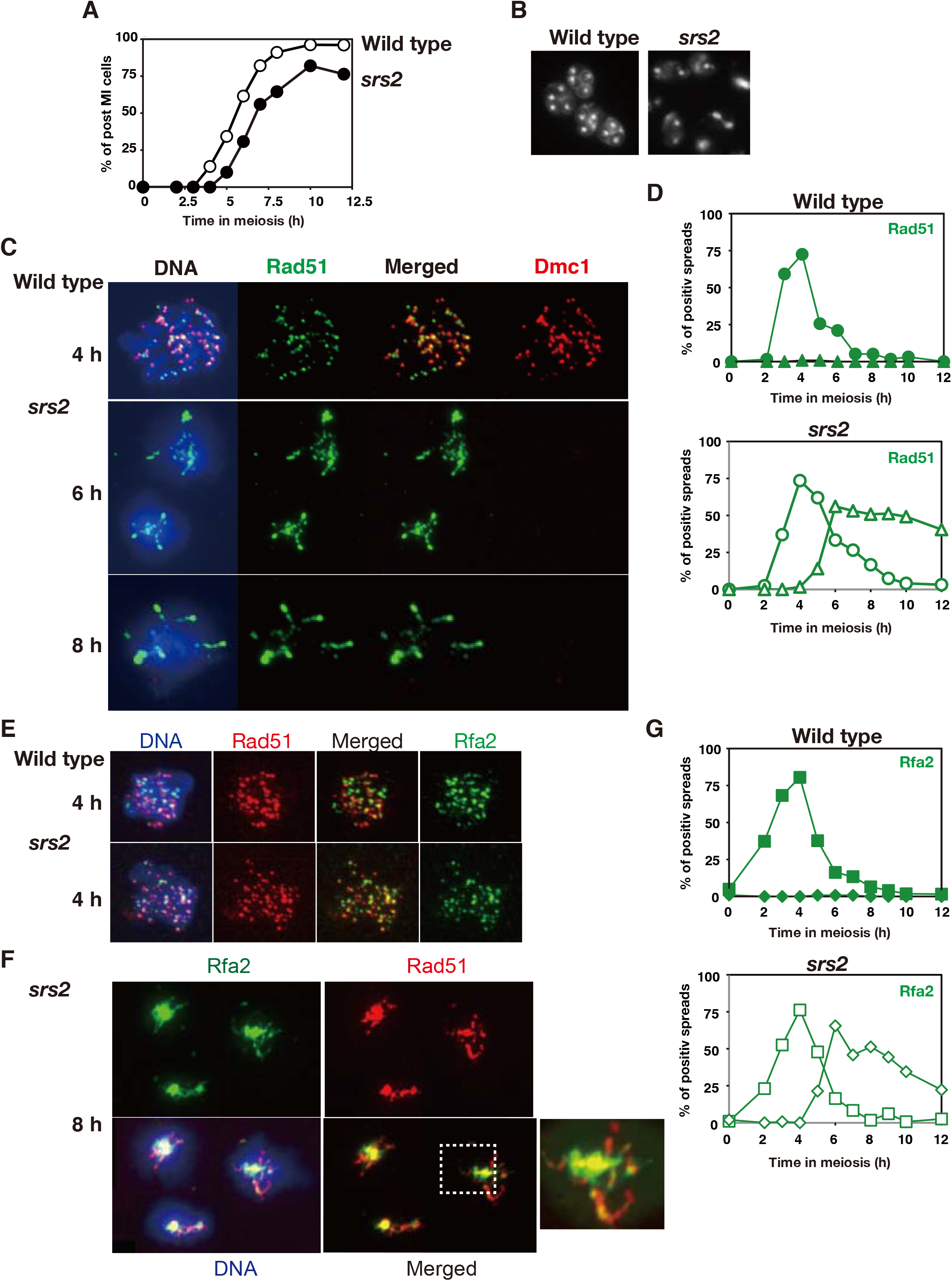
The *srs2* deletion shows Rad51 aggregates during late prophase-I. A. Meiosis I was analyzed by DAPI staining of the wild type (open circles; NKY1303/1543) and *srs2* (filled circles; HSY310/315) cells. The number of DAPI bodies per nucleus was counted in a minimum of 150 DAPI positive cells at each time point. B. DAPI image of wild-type and *srs2* cells at 12 h. C. Immunostaining analysis of Rad51 (green) and Dmc1 (red) on chromosome spreads in wild type (NKY1303/1543) and *srs2* (HSY310/315) cells. Representative images with or without DAPI (blue) dye at 4, 6 and 8 h for wild type and the *srs2* are shown. D. Kinetics of Rad51 focus-positive cells in various yeast strains. A spread with the Rad51 foci is defined as a cell with more than five foci. Spreads containing Rad51 aggregates were also counted. A minimum of 100 cells were analyzed at each time point. Graphs show kinetics of one representative experiment for the wild type cells (top; NKY1303/1543), and *srs2* cells (bottom; HSY310/315). Circles and triangles show spreads with Rad51 foci and aggregate, respectively. E. F. Immunostaining analysis of a component of RPA, Rfa2 (green) with Rad51 (red) in wild type (NKY1303/1543; top) and *srs2* (HSY310/315; bottom) cells at 4 h (E) and at 8 h (F). In F, a dashed square is enlarged in the right. G. Kinetics of Rfa2 foci-positive cells in wild type (NKY1303/1543; top) and *srs2* (HSY310/315; bottom) cells as shown in (B). Circles and triangles show spreads with Rfa2 foci and aggregates, respectively.

### The *srs2Δ* mutant showed a defect in meiotic DSB repair

We analyzed meiotic recombination defects the *srs2* deletion mutant in more detail. First, we checked the repair of meiotic DSBs in the mutant by Southern blotting. DSB formation was monitored at the *HIS4::LEU2* locus, an artificial meiotic recombination hotspot in chromosome *III* (Fig. S1B)(Cao et al. 1990). In wild type, DSB frequencies reached its maximum value at 3 hours of meiosis (~10% of total signals) and then decreased gradually (Fig. S1C, D). The *srs2Δ* accumulates DSB at higher levels (~20%) with more hyper-resection than wild type and delays the disappearance by ~ 2h. (Fig. S1D), indicating that Srs2 is required for efficient meiotic DSB repair. We also checked the formation of two recombinant species, crossover (CO) and non-crossover (NCO) at the same locus. The *srs2Δ* reduces both CO and NCO to 52% and 64% of the wild-type levels (at 6 h; Fig. S1C, D), respectively. These show that Srs2 is necessary for efficient formation of meiotic recombinants. This is consistent with previous return-to-growth experiment showing delayed recombinant prototroph formation in the *srs2Δ* mutant (Palladino and Klein 1992).

During meiotic prophase, homologous chromosomes are tightly coupled with the formation of the synaptonemal complex (SC), a zipper-like chromosome structure linking two homologous chromosomes. Zip1 is a component of the central region of SC, which serves as a marker for synapsis (Sym et al. 1993). A defect in meiotic recombination results in defective SC formation. We checked the SC formation in the *srs2Δ* mutant by immuno-staining analysis of Zip1 on chromosome spreads as well as a meiosis-specific cohesin component, Rec8 (Fig. S1E). We classified three categories according to Zip1 staining; Dotty Zip1 (Class I), partially extended (Class II) and fully-elongated (Class III), which roughly correspond with leptotene, zygotene and pachytene stages, respectively. In wild type, ~66% of nuclei contained full-elongate Zip1 lines at 4 h and Zip1 signal gradually disappeared from chromosomes. In *srs2Δ* strains, although Zip1 focus-positive nuclei exceeded 80% at 4 h, the proportion of cells with fully-elongated Zip1 was significantly reduced to 13 and 26% at 4 and 5 h, respectively (Fig. S1F). Consistent with this, the proportion of polycomplexes (PCs), which are an aggregate of Zip1, dramatically increased; ~60% of the *srs2Δ* nuclei contained PCs at 4 h (Fig. S1G). SCs disassembled more slowly in the mutant than wild type, consistent with delayed meiotic DSB repair (Fig. S1F).

### The *srs2Δ* mutant accumulated aggregates of Rad51 during late meiotic-prophase I

Immuno-staining analysis of chromosome spreads can detect recombination proteins such as Rad51 and Dmc1 on the spreads as a focus, which marks a site of ongoing recombination (Bishop 1994). Previous study indicated that the number of Rad51 foci on chromosome spreads in the *srs2Δ* mutant at 4 h incubation of SPM is slightly reduced compared to those in wild-type (Sasanuma et al. 2013b). We performed kinetic analysis of Rad51 and Dmc1 focus formation. In wild-type cells, dotty signals of both Rad51 and Dmc1 peaked at 4 h of meiosis (Figs. 1C and S2A). The appearance of Rad51 foci in cells lacking Srs2 is slightly delayed, and the disappearance of the foci is delayed relative to wild-type cells (Fig. 1D), consistent with delayed DSB repair in the mutant.

Interestingly, after disappearance of Rad51 foci, we observed reappearance of Rad51 staining with a unique structure after 5 h incubation in the *srs2Δ* mutant (Figs. 1C and S2A). This staining shows clustering of beads-in-line of Rad51 foci, in which 1-5 bright aggregates of Rad51 are connected with each other through thin threads containing Rad51 as well as much simple big aggregation of Rad51 (referred to as Rad51 aggregates) (Fig. 1C). The formation of Rad51 aggregates reach a plateau at 6 h, slightly decreases thereafter, but some cells at 10 or 12 h contained Rad51 aggregates (Fig. 1D), when most of *srs2* mutant cells finished MII (Fig. 1A). At 6 and 12 h, 56 and 40 percent of cells contained aggregates of Rad51, respectively (Fig. 1D, bottom).

Interestingly, this aggregate staining is specific to Rad51, not seen to Dmc1 (Figs. 1C and S2A). Western blots show that Dmc1 and its mediator Mei5 (Hayase et al. 2004) are still present at MI and MII (Fig. S2B). On the other hand, like Rad51 foci, we do see the aggregates of Rad52, a mediator of Rad51 (Shinohara and Ogawa 1998), on chromosomes only in the *srs2Δ* mutant, but not in wild type cells at late times (Fig. S2C, D). We also found that a Rad51 -inhibitor protein, Hed1 (Tsubouchi and Roeder 2006), formed an aggregate with Rad51 with co-localization (Fig. S2E, F). The kinetics of appearance of Rad52 and Hed1 aggregates in the *srs2Δ* mutant are similar to those of Rad51 (Fig. S2D, F).

In order to know the nature of the late Rad51 foci/aggregates, we also studied the localization of RPA (Rfa2, a middle subunit of RPA) at late prophase I of the *srs2Δ* mutant. Immuno-staining showed that, in addition to early RPA foci (Fig. 1E, F), like Rad51-aggregates, aggregate staining of Rfa2 re-appeared at late times of the *srs2* meiosis; e.g. 6-10 h (Fig. 1G). Closer examination reveals that RPA also exhibits a long-line like staining (Fig. 1F). The kinetics of Rfa2 aggregates in the *srs2* mutant is very similar to that of Rad51 (Fig. 1D, G). Some RPA lines and aggregates co-localized with Rad51 lines and aggregates (Fig. 1F). This suggests that the formation of ssDNAs during late prophase-I in *srs2* cells.

One possibility is that Rad51 aggregates bind to DNA damage in ribosomal DNA (rDNA) region, whose segregation defect is often observed in the recombination defective mutants (Li et al. 2014). We co-stained Rad51 with anti-Nop1, a marker for an rDNA region (Schimmang et al. 1989). As shown in Fig. S3A, a single Nop1 signal does not co-localize with late Rad51 aggregates as well as early Rad51 foci in the *srs2* mutant. This excludes the possibility that late Rad51 aggregates are induced by abnormal recombination in the rDNA repeat.

In order to know the relationship of Rad51 aggregate formation with chromosome segregation, we performed whole cell immuno-staining for Rad51 and Dmc1. At early time points, both wild-type and *srs2Δ* mutant cells showed punctate staining for both Rad51 and Dmc1 with some background diffuse staining in a nucleus (Fig. 2A). Rad51-positive nuclei appear at 2 h, peaks at 4 h, and then disappear in wild-type cells while the positive nuclei peaks at 5 h in the *srs2* cells (Fig. 2C). Consistent with results for chromosome spreads (Fig. 1C), the *srs2Δ* cells start to show a big aggregate of Rad51, but not of Dmc1 in nuclei from 5 h and this staining reached to plateau at 8 h (Fig. 2C). Rad51 aggregates in a nucleus often contained thin lines and the number of the aggregate varies up to 2-5 per a nucleus. Importantly, we could also detect Rad51 aggregates in *srs2Δ* cells with two and four big DAPI bodies in a cell, which correspond with cells finishing MI and MII, respectively (Fig. 2B, D). This suggests that DNA damage associated with Rad51 aggregates does not induce delay or arrest of the progression of meiosis. To see the DNA damage checkpoint activation at late meiosis of the *srs2* cells, we analyzed the phosphorylation status of Hop1, which is a substrate of Mec1/ATR and Tel1/ATM as a marker of the activation (Carballo et al. 2008), and found that the *srs2* cells accumulated more phosphorylated Hop1 at 4 h compared to wild type and showed residual phosphorylation during late time points (Fig. 2E).

**Figure 2.**
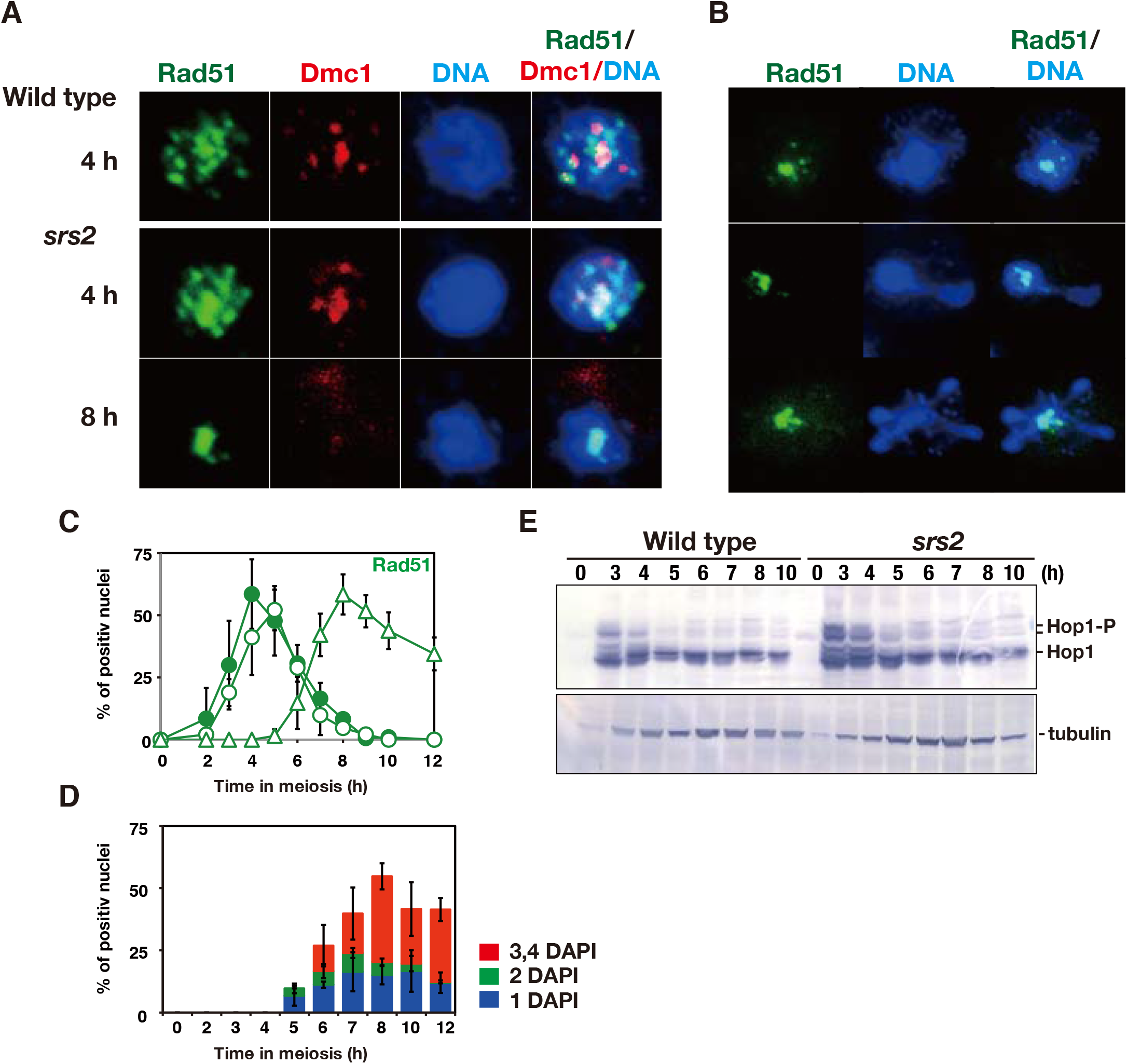
Rad51 aggregates in the *srs2* deletion pass through Meiosis I and II. A, B. Whole cell immunostaining analysis of Rad51 (green) and Dmc1 (red) in wild type (NKY1303/1543) and *srs2* (HSY310/315) cells at 4 and 8 h in meiosis. In (B), cells with Rad51 in prophase-I (top), after MI (middle), and after MII (bottom) are shown. C. Kinetics of Rad51 foci-positive cells in wild-type and *srs2* cells. A foci-positive cell is defined as a cell with more than five foci (closed circles, wild type, NKY1303/1543; open circles, *srs2* cells, HSY310/315). A minimum of 100 cells were analysed at each time point. Graphs show the mean values with S.D. from three independent experiments. D. Kinetics of *srs2* cells containing Rad51 aggregates prior to MI (blue; one DAPI body in a cell) after MI (green; two DAPI bodies in a cell) and MII (red; three or more DAPI bodies in a cell) are shown. Mean values with S.D. from three independent experiments are shown. E. Western blotting analysis of Hop1 phosphorylation during meiosis. Cell lysates at different time points in meiosis in wild type (NKY1303/1543) and *srs2* (HSY310/315) cells were probed with anti-Hop and anti-tubulin antibodies. Phosphorylated Hop1 (shown as “Hop1-P”) shows slower mobility relative to un-phosphorylated Hop1.

### Rad51-aggregate formation in the *srs2* mutant depends on Spo11

To know the nature of Rad51 aggregates in the *srs2Δ* mutant, we looked for genetic requirement of the aggregate formation in the mutant. Rad51–aggregate formation in *srs2Δ* cells is dependent on DSB formation, since a catalytic-dead *spo11* mutation, *spo11-Y135F* (Keeney et al. 1997), almost abolishes both early focus and aggregate of Rad51 staining in the *srs2Δ* mutant (Fig. S3B). It is likely that early DSB-related events in the *srs2Δ* cells may trigger Rad51 aggregates during late meiosis.

In mitosis, the *sgs1* mutation is synthetic lethal with the *srs2* mutation, indicating a redundant role of these two helicases (Gangloff et al. 2000). Sgs1 helicase, together with Top3 and Rmi1, is known to prevent the formation of the untangled chromosomes. The absence of Sgs1 results in abnormal meiosis divisions due to accumulation of un-resolve recombination products involving multi-chromatids (Jessop and Lichten 2008; Jessop et al. 2006; Oh et al. 2007; Oh et al. 2008; Tang et al. 2015). In mammals, the lack of Sgs1 ortholog, BLM helicase, induces anaphase bridges, which are associated with DNA damage generated during S-phase (Biebricher et al. 2013; Chan et al. 2007). We examined the late Rad51 aggregate formation in a meiotic-null allele of *sgs1, sgs1-mn (CLB2p-SGS1)* (Oh et al. 2007). The *sgs1-mn* forms early Rad51 foci with delayed disappearance in prophase of MI, but, unlike the *srs2*, the mutant does not form late Rad51 aggregates (Fig. S3C, D), suggesting that unresolved recombination intermediates formed in the absence of the Sgs1 do not trigger Rad51 aggregates formation.

### The effect of Rad54 depletion on the kinetics of Rad51 aggregates in the *srs2* mutant

We postulated that some Rad51 aggregates turned over during meiosis and could expect to stall its dynamics by blocking late stage of the recombination reaction. We focused on Rad54, which functions at post-assembly stage of Rad51 (Shinohara et al. 1997b), and tried to examine the effect of *RAD54* deletion on Rad51 aggregates. However, it is reported that the *rad54* deletion is synthetically lethal with the *srs2* deletion (Klein 2001; Palladino and Klein 1992; Schild 1995). To circumvent this, we used Rad54-anchor away system, which specifically depletes nuclear Rad54 fused with RFB by the addition of the drug rapamycin (Haruki et al. 2008; Subramanian et al. 2016). The *srs2 RAD54-RFB* cells grow normally in the absence of rapamycin while the *srs2 RAD54-RFB* cells grow poorly on the plate containing the drug, confirming synthetic lethality of the *rad54* and *srs2* (Fig. 3A). In order to know the functional relationship between Rad54 and Srs2 during late meiosis, first, we added rapamycin at 4 h to *RAD54-RFB* and *srs2 RAD54-RFB* cells and analyzed both spore viability and Rad51 foci. The *srs2 RAD54-RFB* decreased spore viability to 64% in the absence of the drug. As reported (Shinohara et al. 1997b), *RAD54-RFB* cells decreased spore viability to 48% in the presence of the drug. Addition of the rapamycin also reduced the spore viability of the *srs2 RAD54-RFB* to 24%, indicating the additive effect of the *srs2* deletion and *RAD54* depletion on spore viability (Fig. 3B). *RAD54* depletion does not affect delayed MI progression in the *srs2* deletion (Fig. 3C). As in wild-type cells, *RAD54-RFB* cells showed normal assembly and disassembly of Rad51 foci in the absence of the drug (Rapa^−^; Fig. 3D, E). However, we found that, from 5 h, one hour after the addition of the drug (Rapa^+^), a new class of Rad51 staining appeared. This class contains 5-10 brighter foci of Rad51, called “Rad51 clump”, which is distinct from the typical Rad51 foci and aggregates (Fig. 3D). This Rad51 clamp peaks at 6 h and then disappears (Fig. 3E), indicating the role of Rad54 in the post Rad51-assembly stage. In the absence of rapamycin, the *srs2 RAD54-RFB* mutant shows the similar kinetics for both Rad51 foci and aggregates to the *srs2Δ* mutant. By the addition of the rapamycin at 4 h, like *RAD54-RFB* cells, the *srs2 RAD54-RFB* mutant formed Rad51 clamp from 5 h and showed the similar kinetics to that in the *RAD54-RFB* (Rapa^+^). In addition, Rad51 aggregates appeared at 5 h and accumulated during further incubation. Rad51 aggregate kinetics in the absence of *RAD54* (Rapa^+^) is delayed relative to its presence (Rapa^−^) (Fig. 3E). This result indicates that Rad51 clumps formed without Rad54 are independent of Srs2. And also, Rad51 aggregate kinetics in the *srs2* cell is independent of Rad54 function.

**Figure 3.**
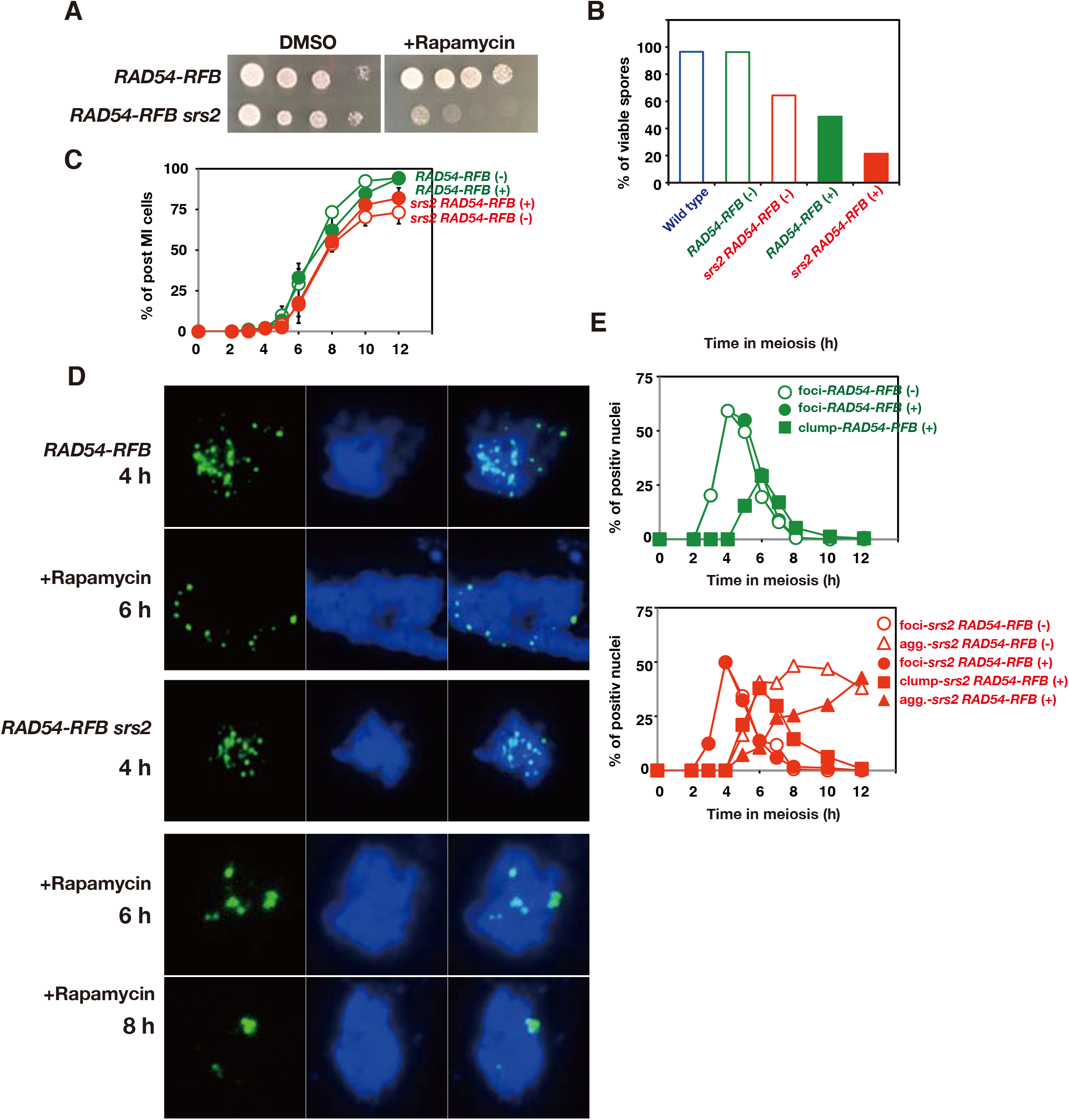
Rad54-depletion induces abnormal Rad51 assembly. A. Mitotic plate assay to confirm synthetic lethality of *srs2* RAD54-anchor on YPAD plates containing 1 μM rapamycin. B. Spore viability of *RAD54-RFB* (H7790/7791) and *RAD54-RFB srs2* (HYS71/82) cells in the absence (−) and the presence (+) of rapamycin. C. Kinetics of MI entry in *RAD54-RFB* (H7790/7791) and *RAD54-RFB srs2* (HYS71/82) cells in the absence (−) and the presence (+) of rapamycin. Rapamycin was added at 4 h at a concentration of 1 μM. Graphs show the mean values with S.D. from three independent experiments. D. Immunostaining analysis of Rad51 (green) and Dmc1 (red) on chromosome spreads in *RAD54-RFB* (H7790/7791) and *RAD54-RFB srs2* (HYS71/82) cells in the absence and the presence of rapamycin. Rapamycin (1 μM) was added at 4 h in meiosis. Representative images of Rad51 staining with or without the addition of rapamycin are shown. E. Kinetics of Rad51 foci-positive cells in *RAD54-RFB* (top green graph; H7790/7791) and *RAD54-RFB srs2* (bottom red graph; HYS71/82) cells in the absence (−) or the presence (+) of rapamycin. Rad51 foci (circles), -clumps (square), and –aggregates (triangles); open symbols (without rapamycin) and closed symbols (addition of the rapamycin at 4 h).

### Rad51 aggregate in the *srs2* mutant is dependent of pachytene exit but is independent of the onset of meiosis I

Rad51 aggregates in the *srs2Δ* mutant are formed at late times during prophase-I. To know the relationship between the focus formation and the progression of meiosis, we first analyzed the Rad51 aggregate formation in the *srs2Δ* with the *ndt80* mutation, which induces pachytene arrest due to the inability to express genes necessary for exit from mid pachytene stage (Xu et al. 1995). Staining of chromosome spreads in *ndt80* cells reveal accumulation of cells with Rad51 foci (Figs. 4A and S3E), which is induced by persistent DSB formation during pachytene arrest by the *ndt80* (Carballo et al. 2013). At later times, the *ndt80* mutant showed the reduced number of Rad51 foci compared to early time points (Fig. S3E). However, Rad51 foci seemed to turn over less efficiently in the *ndt80* mutant (Fig. S3E, F). Little Rad51 aggregate formation was seen in *srs2 ndt80* cells arrested at mid-pachytene both on chromosome spreads and in whole cells (Figs. 4A and S3E). This indicates that the formation of Rad51 aggregates in the *srs2* mutant depends on Ndt80, thus after the exit of mid-pachytene stage.

**Figure 4.**
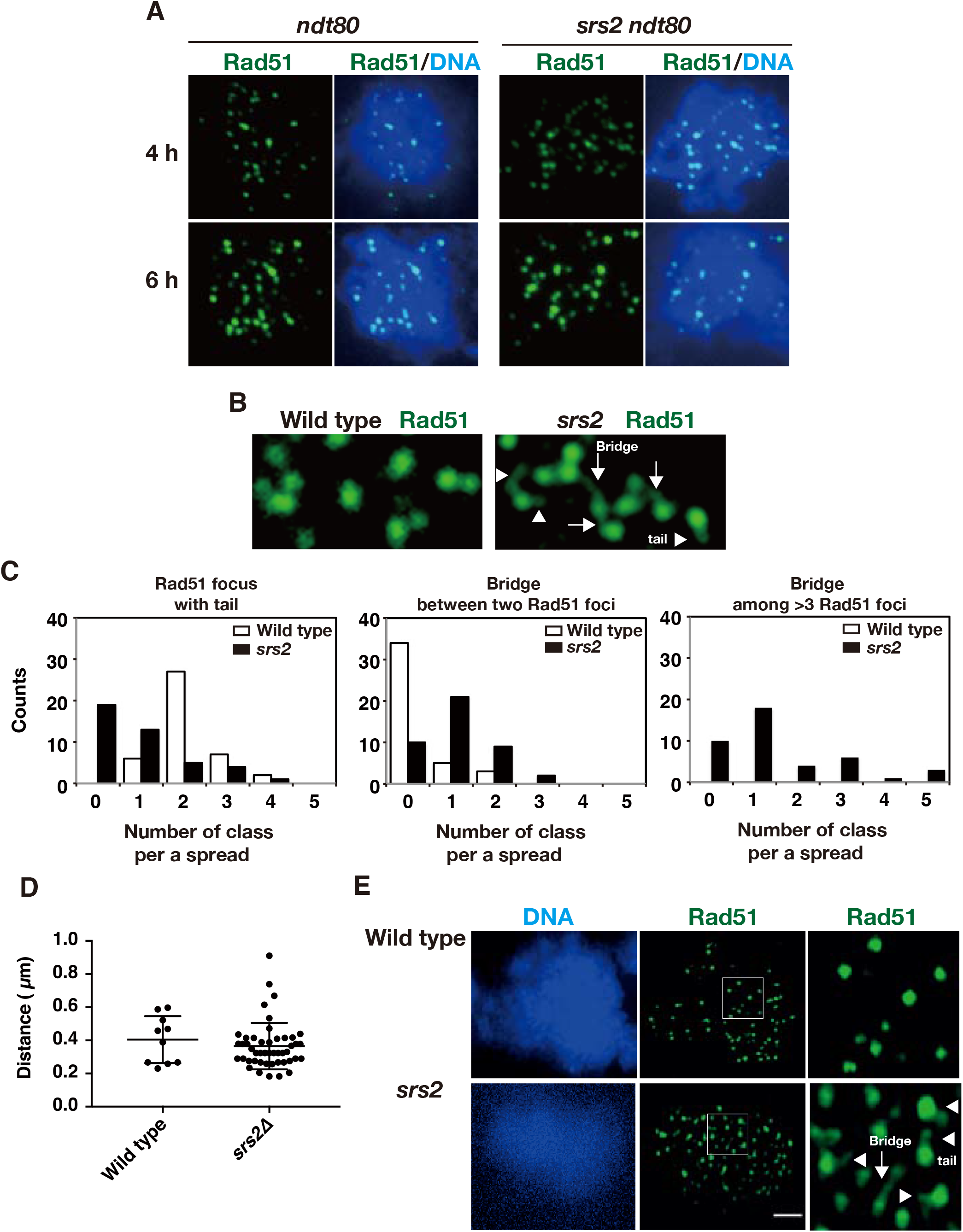
Rad51 aggregate forms in the absence of chromosome segregation. A. Immuno-staining analysis of Rad51 in *ndt80* (HSY596/597) and *srs2 ndt80* (LPY058/059) cells. B. Rad51-bridges in *srs2* cells at 4h. Rad51 tail and bridge are shown in arrowheads and arrows, respectively. C. Rad51-tail or bridge is classified into three classes; Rad51 focus with tail (left), Rad51 bridge between two foci (middle), Rad51 bridge among three or more foci (right). On each spread, the number of each class per a spread was counted, and then a count of the spreads in each class is shown. 42 spreads of wild-type (NKY1303/1543) and *srs2* (HSY310/315) cells were analyzed and counted. D. The length of the Rad51 bridge between two Rad51 foci was measured and plotted. Three horizontal lines from the top indicate the 75, 50 (median), and 25 percentiles, respectively. *P*=0.39; Mann-Whitney *U* test. E. SR-SIM microscopic observation of Rad51 (green) in wild-type (NKY1303/1543) and *srs2* (HSY310/315) cells. Representative image DAPI (blue; left) dye and Rad51 (green, middle) is shown. White insets in middle images are shown in a magnified view at right. The bar indicates 2 μm.

When the kinetics of Rad51 aggregate formation in the *srs2* mutant was compared to kinetics of meiosis I entry, Rad51 aggregate in the *srs2* mutant appear 1 h earlier than the entry into meiosis I (Fig. 1D). To confirm this, we blocked the microtubule dynamics by treating cells with a benomyl, a microtubule depolymerization drug. As shown previously (Hochwagen et al. 2005), the addition of benomyl to yeast meiosis at 4 h prior to the formation of the aggregates, largely suppressed the entry of meiosis I, thus the onset of anaphase-I, in both wild-type and *srs2* cells (Fig. 5A). The treatment with benomyl does not affect Rad51-focus kinetics in both wild-type and *srs2* mutant (Fig. 5A, C). Moreover, the *srs2* cells formed Rad51 aggregates in the presence of benomyl with similar kinetics in its absence (mock treatment with DMSO) (Fig. 5C). This indicates that Rad51-aggregate formation in the *srs2* mutant occurs in the absence of microtubule dynamics, thus chromosome segregation, suggesting that Rad51 aggregate formation in *srs2* mutants is associated with an event during late prophase-I, not with events during the metaphase-I or anaphase-I.

**Figure 5.**
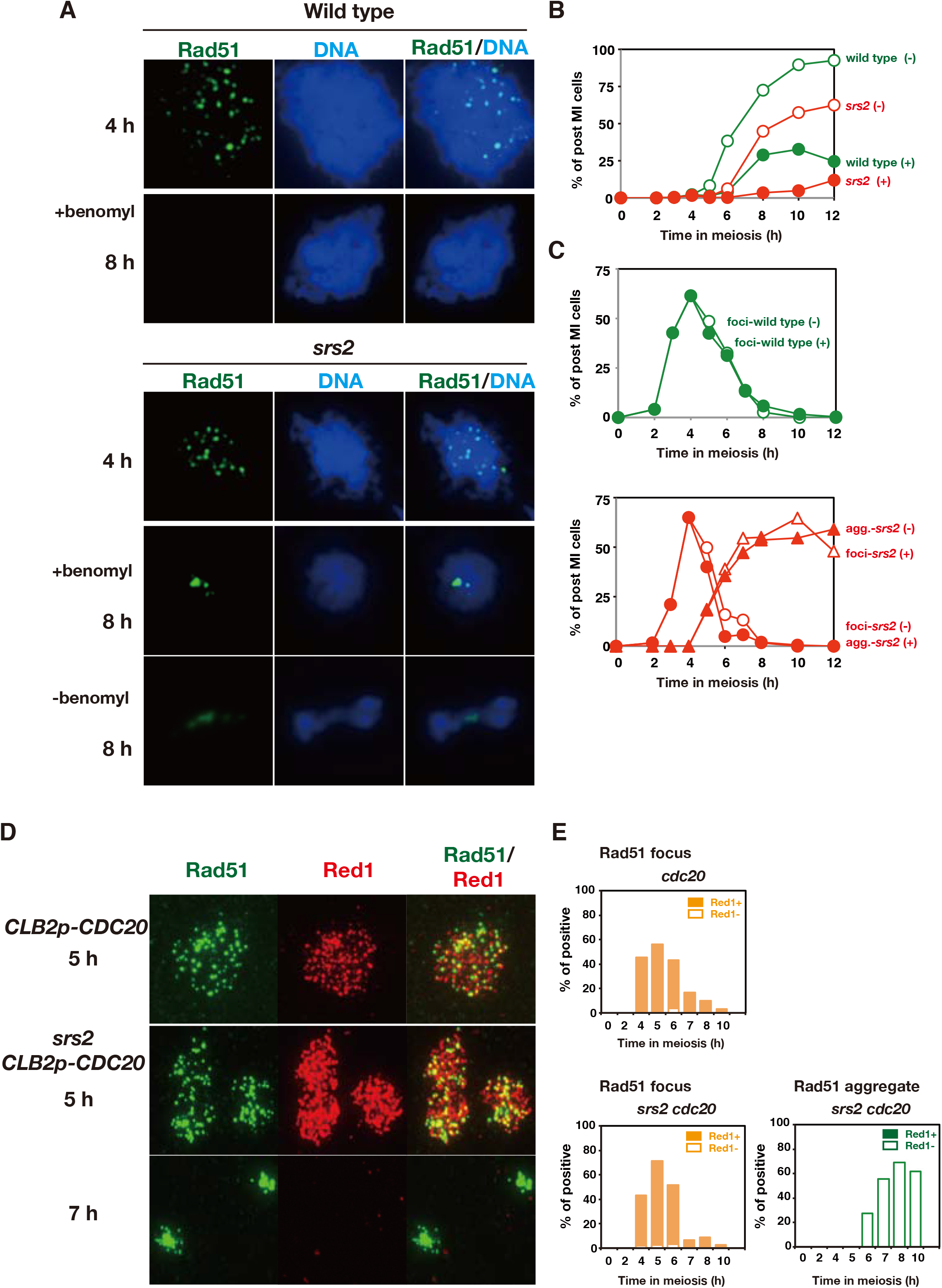
Rad51 aggregate forms in the absence of chromosome segregation. A. Immunostaining analysis of Rad51 (green) in wild type (NKY1303/1543) and *srs2* (HSY310/315) cells in the presence of benomyl. The benomyl was added at 4 h at a concentration of 120 μg/ml. B. Kinetics of MI entry in wild type (green) and *srs2* (red) cells in the absence (open symbols) or the presence (closed symbols) of benomyl. C. Kinetics of Rad51 foci and Rad51 aggregates in wild type (top, green) and *srs2* (bottom, red) cells in the absence (open) or the presence (closed) of benomyl. Circles, Rad51 foci; triangles, Rad51 aggregates. D. Immuno-staining analysis of Rad51 and Red1 in *CDC20-mn* (YFY74/77) and *srs2 CDC20-mn* (YFY80/83) cells. The chromosome spreads at 5 and 7 h were immuno-stained against Rad51 (green) as well as chromosome protein Red1 (red). E. Kinetics of Rad51 aggregate-positive cells in Red1-postive and -negative spreads. Rad51-focus and Rad51-aggregate positive spreads were classified into Red1-negative (open bars) and Red1-positive (closed bars) at each time point. At each time point, more than 50 spreads were counted.

### Rad51 aggregates in the *srs2* mutant appear when SC is disassembled

In order to confirm that Rad51-aggregate formation in the *srs2* is independent of the onset of anaphase I, we used a meiosis-specific null mutant of the *CDC20*, which encodes an activator of Anaphase promoting complex/cyclosome (APC/C), the *cdc20-mn (CLB2p-CDC20)*. As reported previously (Lee and Amon 2003), the *cdc20-mn* shows an arrest at the onset of anaphase I. In the *cdc20-mn*, Rad51 foci appear and disappear like in wild-type control. As expected from the results with benomyl, the Rad51-aggregate formation occurs after the disappearance of Rad51 foci in the *srs2 cdc20-mn* double mutant as in the *srs2* mutant (Fig. 5D, E). This supports the notion that Rad51-aggregate formation in *srs2* mutant is independent of the entry into anaphase-I, thus chromosome segregation.

The relationship between the formation of Rad51 aggregates and late meiotic prophase I such as SC disassembly was compared by immuno-staining of Rad51 with Zip1 (Fig. S4A). After the pachytene exit, the central region of SCs is dismantled as seen in the loss of Zip1 -line signals from chromosomes (Sym et al. 1993). The *srs2* cells containing Rad51 aggregates were almost negative for Zip1 lines (Fig. S4A).

We also performed the staining of Red1, which is a component of chromosome axes (Smith and Roeder 1997). Most cells with Rad51 foci at 3-5 h are almost positive for Red1 staining in both *cdc20-mn* and *srs2 cdc20-mn* cells (Fig. 5D, E). In contrast, *srs2 cdc20-mn* cells with Rad51 aggregates were negative for Red1 signal. These indicate that Rad51 aggregate formation in the *srs2* occurs after or during disassembly of Red1-axes. This is confirmed in the background of wild type too (Fig. S4B, C).

We confirmed this by staining of Rec8, a kleisin subunit of cohesin (Klein et al. 1999). At late time points such as 6 h, Rec8 showed dotty staining compared to 5 h (Challa et al. 2019), when most of Rec8 show line staining. Rec8 line positive spreads contained Rad51 foci (Fig. S4D, E). In *srs2* cells with or without *cdc20-mn*, Rad51 aggregates are predominantly seen in cells with Rec8-dots (Fig. S4D, E).

### The *srs2* mutant accumulated bridge staining of Rad51 between two recombination foci during early prophase-I

During our staining analysis, we noticed that the *srs2* cells show very unique thin line staining of Rad51 during early prophase such as 4 h (Fig. 4B). The thin Rad51-line in *srs2* cells is connected from one Rad51 focus to the other focus/foci, which we refer to as “Rad51 bridge”. At least one clear Rad51 bridge between two Rad51 foci were observed at ~40% frequency of *srs2* spreads at 4 h (middle graph of Fig. 4C). A few Rad51 bridges were seen in wild type. We also found the Rad51-bridge staining among more than three Rad51 foci in *srs2* cells, but not in wild type (right graph of Fig. 4C). Careful examination of Rad51 foci in the wild type often detected a Rad51 focus with “single tail (or whisker)” (left graph of Figure 4C). The number of Rad51 tail from a single Rad51 focus is almost one. There is few focus with more than 2 tails. When measured the length of the bridge between two foci, we found both wild-type and the *srs2* cells show similar distribution of the lengths (Fig. 4D). These results indicate that Srs2 suppresses the formation of Rad51 bridges. Indeed, the *srs2* cells increased the frequency of the Rad51 bridges and more connections among more than two foci relative to the wild type (Fig. 4C).

We then used super-resolution microscopy to analyze Rad51 localization on meiotic chromosomes at high resolution. A structural illumination microscope (SIM) was used to determine Rad51 localization in wild type and *srs2* cells at 4h (Fig 4E). As shown above, in the *srs2* mutant, we detected both Rad51 bridges and tails more than in wild type. The wild type the *srs2* mutant shows Rad51 foci with tail/bridge at a frequency of 15.4±4.2% (n=18) and 54.3±8.5% (n=20), respectively.

The average length of the bridge is ~0.4 μm (Fig. 4D). If the bridge is postulated to consist of a single Rad51 filament on the ssDNA, which is extended 2-fold relative to the B-form DNA (Ogawa et al. 1993), we can calculate the bridge contains ~600 nt (400 nm/2X3.3/10.5). This might be a range of reasonable estimate ssDNA length at a single DSB site with ~900 nt (Mimitou et al. 2017; Zakharyevich et al. 2010). Rad51 bridge described here might be similar to the staining of “ultra fine bridge” seen in anaphase of damage mammalian cells (Chan and Hickson 2011) (see Discussion). The formation of Rad51 thin bridges in early prophase-I of the *srs2* cells suggests entanglement of recombination intermediates.

### Little chromosome breaks are formed in the *srs2* mutant during late prophase-I

In order to detect chromosomal breaks, we tried to analyze chromosome status by pulse field gel electrophoresis (PFGE). At 3, 4 h time points in both wild-type and *srs2* mutant cells, chromosomal band becomes a smear due to the introduction of DSBs (Fig. S5). While the smear pattern disappeared at 5 h in wild type, the *srs2* mutant showed persistent smear bands by 5 h and then disappeared. Interestingly, we again detected smear bands at 10 and 12 h when most of the *srs2* diploid makes spores, indicating the formation of DSBs in the *srs2* spores. The smear bands were barely observed in wild type spore. More importantly, the *srs2* cells did not show breaks at 5-8 h, when Rad51 aggregates are induced.

## Discussion

### Rad51 bridges and Rad51 aggregates in the *srs2* mutant

The *srs2Δ* mutant shows decreased levels of CO and NCO relative to the wild-type, indicating a positive role of Srs2 in meiotic recombination (pro-recombination role). This weak defect in the recombination is consistent with delayed DSB repair (delayed disassembly of Rad51 foci) as well as defective SC formation in the mutant. Our studies also showed that the *srs2* mutant is partially defective in a step after the DSB processing. However, this “weak” defect in the recombination cannot explain reduced spore viability of the mutant, since the mutants with 50% reduction of CO show high spore viability; e.g. *spo11, xrs2, msh4/5* hypomorphic mutants (Martini et al. 2006; Nishant et al. 2010; Shima et al. 2005). Consistent with low spore viability of the mutant, we and others detected abnormal chromosome segregation in *srs2* meiosis, suggesting the presence of DNA abnormality in the mutant.

In this study, we described “unusual” DNA damage formed in the absence of Srs2 helicase during meiosis. This damage is marked with the association of the recombination protein, Rad51, with a large quantity, which we refer to as “Rad51 aggregate”. The Rad51 aggregate is not a protein aggregate since it contains another recombination protein, Rad52, as well as RPA, but not meiosis-specific recombination proteins such as Dmc1. The presence of RPA strongly suggests the presence of ssDNAs. Indeed, thin line-like staining of Rad51 and RPA emanating from the aggregate are often observed.

The formation of Rad51 aggregates in *srs2Δ* mutant requires Spo11 catalytic activity, thus DSB formation. On the other hand, kinetic analysis revealed that Rad51 aggregates in *srs2Δ* mutant appear in late prophase-I after the disappearance of Spo11-dependent Rad51 foci associated with meiotic recombination. Rad51 aggregates appear just after the disappearance of “normal” Rad51 foci. This suggests that the formation of Rad51 aggregates occur after the completion of DSB repair such as Rad51-mediated strand invasion. Consistent with this, the *ndt80* mutation, which induces an arrest at mid-pachytene stage, blocks the aggregate formation in the *srs2Δ* mutant. The *ndt80* mutant accumulates dHJ as a product of completion of Rad51-dependent strand invasion (Allers and Lichten 2001), and also shows persistent formation of Spo11-dependent meiotic DSBs (Carballo et al. 2013). Therefore, persistent DSBs and dHJs are unlikely to be directly linked with Rad51 aggregate formation.

Mutant analysis shows the formation of Rad51 aggregates in the *srs2Δ* requires pachytene-exit, but occurs prior to the transition of metaphase-I to anaphase-I, chromosome segregation. Indeed, Rad51 aggregate formation occurs even when chromosome segregation was inhibited by the treatment with a microtubule polymerization inhibitor and the *CDC20* depletion, which delays and blocks the onset of anaphase-I. These indicate that the aggregate formation is induced around the disassembly of meiotic chromosome structure; e.g. diplotene or diakinesis.

One possibility to explain Rad51 aggregate formation in the *srs2* mutant is that, after the exit of Ndt80-execution point, there might be unrepaired DSBs, which could be repaired by Rad51-dependent pathway (but not Dmc1-pathway) during late prophase-I. The *srs2* mutant might be specifically defective in this DSB repair after the pachytene exit. In this pathway, Srs2 may be essential for Rad51 removal, which may lead to the accumulation of unrepaired ssDNAs. However, this is unlikely since even DSB ends formed during pachytene are bound by Dmc1 as well as Rad51. However, the Rad51 aggregates in the *srs2* mutant do not contain Dmc1 even when Dmc1 protein is present in a cell.

Alternatively, Rad51 aggregates and/or its associated DNA damage are formed in two-step process. Frist, DSB repair in the absence of Srs2 may result in the formation of aberrant recombination products/intermediates such as entangled duplexes DNAs (see Fig. 6B). Second, this aberrant product/intermediate might be converted into DNA damage with Rad51 aggregates in late prophase-I. Consistent with this two-step model, we found a novel structure called Rad51-bridge (or whisker), thin lines of Rad51 which connect Rad51 foci. This bridge is seen at early prophase I of the *srs2* mutant more frequently than in wild type.

**Figure 6.**
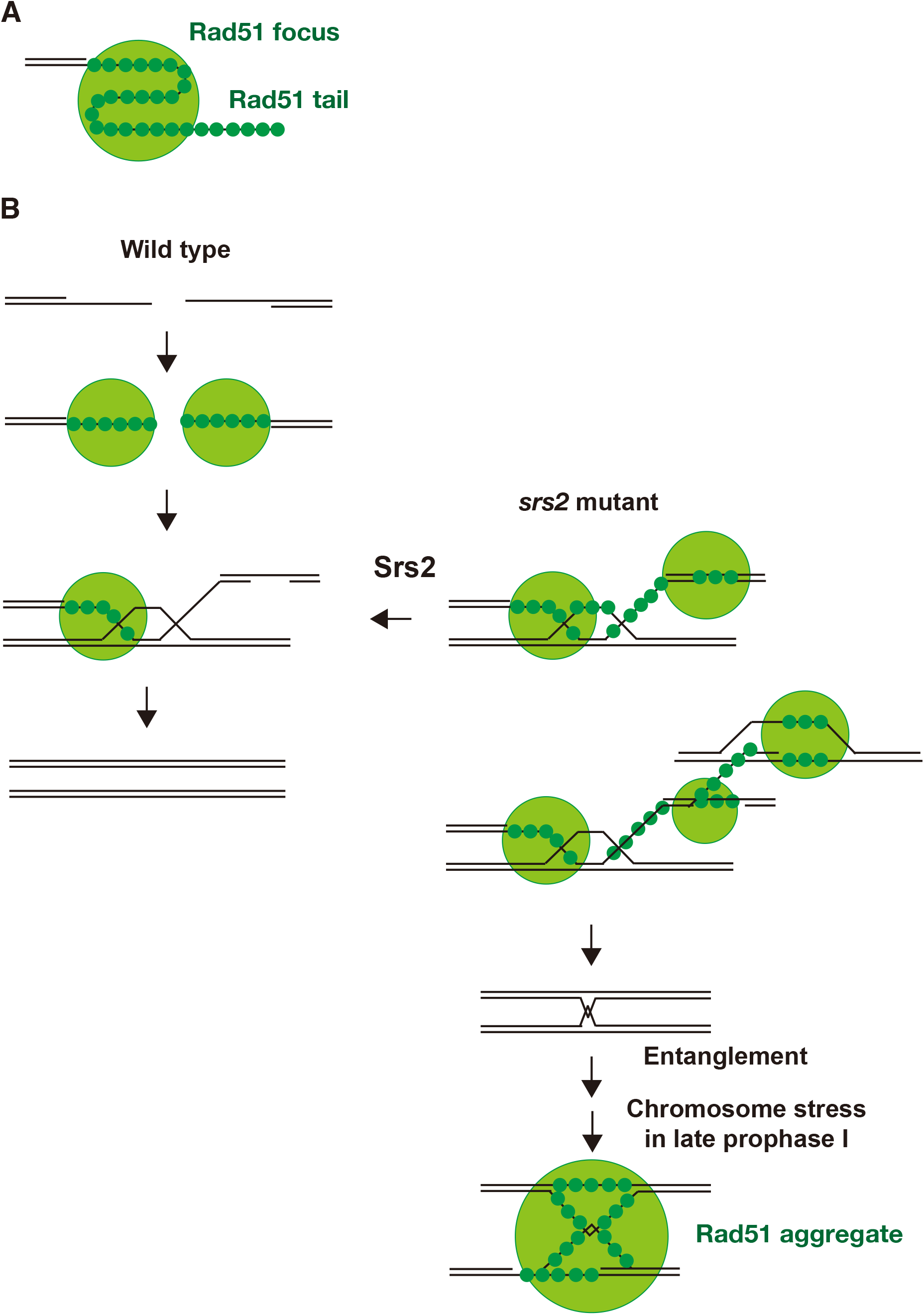
A schematic model of Rad51 bridge and Rad51 aggregate formation in wild type and *srs2* cells. A. A model of Rad51 focus comprised of the Rad51 filament. Rad51 filament is accommodated into a three-dimensional structure. B. A recombination pathway with Rad51 focus and filaments is described. In left, the second end capture by a displaced strand from D-loop, which is mediated by the Rad51 filament may trigger the disassembly of Rad51 filament (focus) on the second end, which is promoted by Srs2. Srs2 also prevents the formation of a multi-invasion intermediate. These abnormal recombination intermediates are processed into a final product, which may contain a pathological damage that may lead to the formation of a long ssDNAs with Rad51 aggregates during late prophase-I.

The presence of Rad51-bridge and -whisker from Rad51 focus suggests that Rad51 focus is not a simple Rad51 filament, rather may contain a three-dimensional configuration of Rad51 filament (Fig. 6A). Rad51-bridge line staining is reminiscent of anaphase bridge or ultra-fine bridge of chromosomes in mammalian cells (Chan et al. 2007). The formation of anaphase bridges in mammalian cells is a two-step process. Although these bridges are formed during M phase with onset of anaphase, the initiation event leading to the bridge formation occurs during S-phase. These bridges are induced by the treatment of the cell with DNA replication inhibitor(s) or in the absence of DNA repair protein such as BLM helicase. The bridges are decorated with repair proteins such as BLM and RPA, but not Rad51.

Rad51 aggregate formation in *srs2* meiosis is clearly different from the anaphase bridge in the following two aspects. First, the formation is not required for chromosome segregation. Second, the initiation event should be Spo11-dependent DSB formation in early prophase-I (meiotic G_2_ phase). If the two-step model as described above is true, the conversion of the aberrant product/intermediate into the aggregate in *srs2Δ* mutant should occur after mid-pachytene exit. Given drastic chromosome morphogenesis such as chromosome compaction and disassembly of the meiosis-specific chromosome structure, SC, occur during late meiotic prophase-I (Challa et al. 2019), these events might induce the aberrant DNA damage with Rad51 aggregates.

Given that Rad51-bridge in the *srs2* mutant is formed between Rad51 foci, Srs2 might play a role of this kind of Rad51-associated DNA entanglement between the two DSB sites (Fig. 6B). One likely intermediate is multiple invasion (Piazza et al. 2017), which are formed by Rad51-mediated strand invasion into multiple loci. Therefore, Srs2 might play a role in resolution of multiple-invasion by controlling Rad51 filament dynamics using its Rad51-dismantling activity.

Bishop and his colleagues show a pair of Rad51 foci during early meiotic pro-phase I are formed in the two ends of a single DSB site (Brown et al. 2015). Thus, it is likely that the Rad51 bridge we observed is formed between a pair of Rad51 foci on the two DSB ends. If so, one likely possibility is that the bridge is a ssDNA between two DSB ends. One way to connect the two DSB ends is bridged by the annealing of ejected ssDNA from the D-loop after the DNA synthesis (Fig. 6B). Since the bridge is mainly seen in the absence of Srs2, we propose that Rad51 dismantling activity of Srs2 promotes the removal of Rad51 from the rejected ssDNA. Moreover, it is likely that Srs2 also remove Rad51 in the other end of the DSB during the second end-capture. This idea could explain the formation of the bridge between two Rad51 foci in the *srs2*, but not in wild type. In wild-type cells, Srs2 seems to remove Rad51 assembly from the intermediates for the second end capture. Importantly, genetic analysis of mitotic recombination in the *srs2* mutant suggest the role of Srs2 to facilitate the annealing of the newly synthesized strand to second resected ends by removing Rad51 from the second end (Elango et al. 2017; Ira et al. 2003; Liu et al. 2017; Mitchel et al. 2013).

We still cannot figure out recombination products formed in the absence of Srs2, which trigger the formation of the Rad51 aggregate. 2D gel analysis has shown that there is few accumulation of abnormal recombination intermediate such as multiple dHJs (Lichten/Goldman, accompanying paper). Thus, multiple dHJs is unlikely. Rather, there might be an entanglement of DNA strands after the completion of the meiotic recombination (Fig. 6B). This intermediate seems to be related to a lethal recombination intermediate formed in the *srs2* mutant during mitosis.

In either scenario, our analysis reveals a novel pathway to protect meiotic cells in late prophase-I from the formation of aberrant DNA damage induced by Spo11. This repair pathway heavily depends on Srs2 function. For this function, Srs2 is almost essential for meiosis. We would like to point out that Rad51 aggregates in the *srs2* mutant is related to lethal recombination intermediates in mitotic cells with *SRS2* deletion, which is postulated to form through two-step model.

Rad51 aggregate-associated DNA damage seems unrepaired during meiosis. During late prophase-I, there should be sister chromatid or other recombination partners to repair the damage, this might be due to the presence of Rad51-inhibitor Hed1, which clearly suppresses Rad51-mediated DNA repair.

The result that CHEF-Southern for chromosome III did not detect any DNA fragmentation at times of Rad51-aggregates in the *srs2Δ* mutant; e.g. 6 and 8 h, implies Rad51 aggregate-associated DNA damage does not contain DSBs. One simple interpretation is that Rad51 aggregates are on either the ssDNA gaps or unwound duplex DNAs.

### No activation of DNA damage checkpoint during late prophase I, meiosis I and meiosis II

In the absence of Srs2, DNA damage with Rad51 aggregates is formed and passed into MI and MII with activation of DNA damage checkpoint, which leads more catastrophic events such as chromosome fragmentation with DSB formation in spores. This may explain quite a big reduction of spore viability of the *srs2* diploid with reasonable levels of meiotic recombination.

The absence of DNA damage-induced delay in late meiosis-I in the *srs2* cells is quite different from cell cycle delay induced by the recombination (pachytene) checkpoint during early prophase-I (MacQueen and Hochwagen 2011; Tsubouchi et al. 2018). In the recombination checkpoint, DSBs and associated ssDNA activate sensor kinases Tel1(ATM) and Mec1(ATR), respectively. During meiosis, activated Mec1 and Tel1 induce the activation of a meiosis-specific kinase, Mek1, Rad53 homolog, by phosphorylating its partner protein Hop1. High Mek1 activity down-regulates Ndt80 activity, thus, blocking the exit of mid-pachytene stage. During meiosis, the activation of mitotic DNA damage downstream kinases, Rad53 and Chk1, is blocked through an unknown mechanism. At late times in the *srs2* mutant, we did not see prolonged phosphorylation of Hop1, thus little activation of Mec1 (and Tel1). This strongly suggests that DNA damage with Rad51 aggregates in the *srs2* is masked by the checkpoint activation or there is no such mechanism in late G2 phase of meiotic cells. Alternatively, although not exclusive with the above, Srs2 may function in the activation of the checkpoint during this phase.

### Role of Rad54 in late recombination

Upon Rad54 depletion after the assembly of Rad51 on the ssDNA during meiosis, we found a novel staining of Rad51 called Rad51 clump, which is different from typical Rad51 foci in wild-type and aggregates in the *srs2* cells. The presence of Rad51 clumps supports the idea of a role of Rad54 after the assembly of Rad51 filaments. Moreover, the Rad54-Rad51, not with Dmc1, may function in the repair of DSBs in late prophase-I. Previous cytological analysis of the *rad54* deletion does not show the formation of Rad51 clump (Shinohara et al. 2000; Shinohara et al. 1997b). One possibility is that Rad54 depletion may remove Rad54-associated proteins also from the nuclei. As a result, we do see clear defect in Rad51 dynamics upon Rad54 depletion in late prophase-I.

In the accompanying paper, Goldman and his colleagues described the formation of Rad51 aggregates during *srs2* meiosis.

## Acknowledgements

We are grateful for Drs. Alstair Goldman and Michael Lichten for sharing unpublished results prior to publication. We thank Dr. Neil Hunter (UC, Davis) for *pCLB2-SGS1* yeast and Dr. Andreas Hochwagen (New York University) for the anchor-away yeast strains. We thank Ms. H. Matsumoto, C. Watanabe, and H. Wakabayashi for excellent technical assistance.

## Author’s contribution

H.S., M.S., and A.S. designed the experiments. H.S., H.S.M.S., Y.F., and L.P. performed all experiments. M.S. provided reagents. H.S., H.S.M.S., and A.S. analyzed the data. A.S. prepared manuscripts with help by H.S., H.S.M.S. and M.S.

## References

Allers T, Lichten M (2001) Differential timing and control of noncrossover and crossover recombination during meiosis Cell 106:47–57

Bani Ismail M, Shinohara M, Shinohara A (2014) Dot1-dependent histone H3K79 methylation promotes the formation of meiotic double-strand breaks in the absence of histone H3K4 methylation in budding yeast PloS one 9:e96648 doi:10.1371/journal.pone.0096648

Biebricher A et al. (2013) PICH: a DNA translocase specially adapted for processing anaphase bridge DNA Mol Cell 51:691–701 doi:10.1016/j.molcel.2013.07.016

Bishop DK (1994) RecA homologs Dmc1 and Rad51 interact to form multiple nuclear complexes prior to meiotic chromosome synapsis Cell 79:1081–1092

Bishop DK, Park D, Xu L, Kleckner N (1992) DMC1: a meiosis-specific yeast homolog of E. coli recA required for recombination, synaptonemal complex formation, and cell cycle progression Cell 69:439–456

Brown MS, Grubb J, Zhang A, Rust MJ, Bishop DK (2015) Small Rad51 and Dmc1 Complexes Often Co-occupy Both Ends of a Meiotic DNA Double Strand Break PLoS genetics 11:e1005653 doi:10.1371/journal.pgen.1005653

Cahoon CK, Hawley RS (2016) Regulating the construction and demolition of the synaptonemal complex Nature Structural & Molecular Biology 23:369–377 doi:10.1038/nsmb.3208

Cao L, Alani E, Kleckner N (1990) A pathway for generation and processing of double-strand breaks during meiotic recombination in S. cerevisiae Cell 61:1089–1101

Carballo JA, Johnson AL, Sedgwick SG, Cha RS (2008) Phosphorylation of the axial element protein Hop1 by Mec1/Tel1 ensures meiotic interhomolog recombination Cell 132:758–770 doi:10.1016/j.cell.2008.01.035

Carballo JA et al. (2013) Budding yeast ATM/ATR control meiotic double-strand break (DSB) levels by down-regulating Rec114, an essential component of the DSB-machinery PLoS genetics 9:e1003545 doi:10.1371/journal.pgen. 1003545

Cejka P, Plank JL, Bachrati CZ, Hickson ID, Kowalczykowski SC (2010) Rmi1 stimulates decatenation of double Holliday junctions during dissolution by Sgs1-Top3 Nature Structural & Molecular Biology 17:1377–1382 doi:10.1038/nsmb.1919

Challa K, Fajish VG, Shinohara M, Klein F, Gasser SM, Shinohara A (2019) Meiosis-specific prophase-like pathway controls cleavage-independent release of cohesin by Wapl phosphorylation PLoS Genetics 15:e1007851 doi:10.1371/journal.pgen.1007851

Chan KL, Hickson ID (2011) New insights into the formation and resolution of ultra-fine anaphase bridges Seminars in cell & developmental biology 22:906–912 doi:10.1016/j.semcdb.2011.07.001

Chan KL, North PS, Hickson ID (2007) BLM is required for faithful chromosome segregation and its localization defines a class of ultrafine anaphase bridges Embo j 26:3397–3409 doi:10.1038/sj.emboj.7601777

Cloud V, Chan YL, Grubb J, Budke B, Bishop DK (2012) Rad51 is an accessory factor for Dmc1-mediated joint molecule formation during meiosis Science 337:1222–1225 doi:10.1126/science.1219379

Elango R et al. (2017) Break-induced replication promotes formation of lethal joint molecules dissolved by Srs2 Nature communications 8:1790 doi:10.1038/s41467-017-01987-2

Gangloff S, Soustelle C, Fabre F (2000) Homologous recombination is responsible for cell death in the absence of the Sgs1 and Srs2 helicases Nat Genet 25:192–194

Haruki H, Nishikawa J, Laemmli UK (2008) The anchor-away technique: rapid, conditional establishment of yeast mutant phenotypes Mol Cell 31:925–932 doi:10.1016/j.molcel.2008.07.020

Hayase A, Takagi M, Miyazaki T, Oshiumi H, Shinohara M, Shinohara A (2004) A protein complex containing Mei5 and Sae3 promotes the assembly of the meiosis-specific RecA homolog Dmc1 Cell 119:927–940

Hochwagen A et al. (2005) Novel response to microtubule perturbation in meiosis Mol Cell Biol 25:4767–4781 doi:10.1128/mcb.25.11.4767-4781.2005

Hong EL, Shinohara A, Bishop DK (2001) Saccharomyces cerevisiae Dmc1 protein promotes renaturation of single-strand DNA (ssDNA) and assimilation of ssDNA into homologous super-coiled duplex DNA The Journal of biological chemistry 276:41906–41912

Hunter N, Kleckner N (2001) The single-end invasion: an asymmetric intermediate at the double-strand break to double-holliday junction transition of meiotic recombination Cell 106:59–70

Ira G, Malkova A, Liberi G, Foiani M, Haber JE (2003) Srs2 and Sgs1-Top3 suppress crossovers during double-strand break repair in yeast Cell 115:401–411

Jessop L, Lichten M (2008) Mus81/Mms4 endonuclease and Sgs1 helicase collaborate to ensure proper recombination intermediate metabolism during meiosis Mol Cell 31:313–323 doi:10.1016/j.molcel.2008.05.021

Jessop L, Rockmill B, Roeder GS, Lichten M (2006) Meiotic chromosome synapsis-promoting proteins antagonize the anti-crossover activity of sgs1 PLoS genetics 2:e155 doi:10.1371/journal.pgen.0020155

Keeney S, Giroux CN, Kleckner N (1997) Meiosis-specific DNA double-strand breaks are catalyzed by Spo11, a member of a widely conserved protein family Cell 88:375–384

Klein F, Mahr P, Galova M, Buonomo SB, Michaelis C, Nairz K, Nasmyth K (1999) A central role for cohesins in sister chromatid cohesion, formation of axial elements, and recombination during yeast meiosis Cell 98:91–103 doi:10.1016/S0092-8674(00)80609-1

Klein HL (2001) Mutations in recombinational repair and in checkpoint control genes suppress the lethal combination of srs2Delta with other DNA repair genes in Saccharomyces cerevisiae Genetics 157:557–565

Krejci L et al. (2003) DNA helicase Srs2 disrupts the Rad51 presynaptic filament Nature 423:305–309

Lee BH, Amon A (2003) Role of Polo-like kinase CDC5 in programming meiosis I chromosome segregation Science 300:482–486 doi:10.1126/science.1081846

Li P, Jin H, Yu HG (2014) Condensin suppresses recombination and regulates double-strand break processing at the repetitive ribosomal DNA array to ensure proper chromosome segregation during meiosis in budding yeast Mol Biol Cell 25:2934–2947 doi:10.1091/mbc.E14-05-0957

Liu J et al. (2017) Srs2 promotes synthesis-dependent strand annealing by disrupting DNA polymerase delta-extending D-loops eLife 6 doi:10.7554/eLife.22195

MacQueen AJ, Hochwagen A (2011) Checkpoint mechanisms: the puppet masters of meiotic prophase Trends in cell biology 21:393–400 doi:10.1016/j.tcb.2011.03.004

Marini V, Krejci L (2010) Srs2: the “Odd-Job Man” in DNA repair DNA Repair (Amst) 9:268–275

Martini E, Diaz RL, Hunter N, Keeney S (2006) Crossover homeostasis in yeast meiosis Cell 126:285–295 doi:10.1016/j.cell.2006.05.044

Mimitou EP, Yamada S, Keeney S (2017) A global view of meiotic double-strand break end resection Science 355:40–45 doi:10.1126/science.aak9704

Mitchel K, Lehner K, Jinks-Robertson S (2013) Heteroduplex DNA position defines the roles of the Sgs1, Srs2, and Mph1 helicases in promoting distinct recombination outcomes PLoS genetics 9:e1003340 doi:10.1371/journal.pgen.1003340

New JH, Sugiyama T, Zaitseva E, Kowalczykowski SC (1998) Rad52 protein stimulates DNA strand exchange by Rad51 and replication protein A Nature 391:407–410 doi:10.1038/34950

Nishant KT, Chen C, Shinohara M, Shinohara A, Alani E (2010) Genetic analysis of baker’s yeast Msh4-Msh5 reveals a threshold crossover level for meiotic viability PLoS genetics 6 doi:10.1371/journal.pgen.1001083

Ogawa T, Yu X, Shinohara A, Egelman EH (1993) Similarity of the yeast RAD51 filament to the bacterial RecA filament Science 259:1896–1899

Oh SD, Lao JP, Hwang PY, Taylor AF, Smith GR, Hunter N (2007) BLM ortholog, Sgs1, prevents aberrant crossing-over by suppressing formation of multichromatid joint molecules Cell 130:259–272 doi:10.1016/j.cell.2007.05.035

Oh SD, Lao JP, Taylor AF, Smith GR, Hunter N (2008) RecQ helicase, Sgs1, and XPF family endonuclease, Mus81-Mms4, resolve aberrant joint molecules during meiotic recombination Mol Cell 31:324–336 doi: 10.1016/j.molcel.2008.07.006

Palladino F, Klein HL (1992) Analysis of mitotic and meiotic defects in Saccharomyces cerevisiae SRS2 DNA helicase mutants Genetics 132:23–37

Petronczki M, Siomos MF, Nasmyth K (2003) Un menage a quatre: the molecular biology of chromosome segregation in meiosis Cell 112:423–440

Piazza A, Wright WD, Heyer WD (2017) Multi-invasions Are Recombination Byproducts that Induce Chromosomal Rearrangements Cell 170:760–773.e715 doi:10.1016/j.cell.2017.06.052

Rong L, Palladino F, Aguilera A, Klein HL (1991) The hyper-gene conversion hpr5-1 mutation of Saccharomyces cerevisiae is an allele of the SRS2/RADH gene Genetics 127:75–85

Sasanuma H, Furihata Y, Shinohara M, Shinohara A (2013a) Remodeling of the Rad51 DNA strand-exchange protein by the Srs2 helicase Genetics 194:859–872 doi:10.1534/genetics.113.150615

Sasanuma H et al. (2008) Cdc7-dependent phosphorylation of Mer2 facilitates initiation of yeast meiotic recombination Genes Dev 22:398–410 doi:10.1101/gad.1626608

Sasanuma H et al. (2013b) A new protein complex promoting the assembly of Rad51 filaments Nature communications 4:1676 doi:10.1038/ncomms2678

Schild D (1995) Suppression of a new allele of the yeast RAD52 gene by overexpression of RAD51, mutations in srs2 and ccr4, or mating-type heterozygosity Genetics 140:115–127

Schimmang T, Tollervey D, Kern H, Frank R, Hurt EC (1989) A yeast nucleolar protein related to mammalian fibrillarin is associated with small nucleolar RNA and is essential for viability Embo j 8:4015–4024

Schwacha A, Kleckner N (1994) Identification of joint molecules that form frequently between homologs but rarely between sister chromatids during yeast meiosis Cell 76:51–63

Shima H, Suzuki M, Shinohara M (2005) Isolation and characterization of novel xrs2 mutations in Saccharomyces cerevisiae Genetics 170:71–85 doi:10.1534/genetics.104.037580

Shinohara A, Gasior S, Ogawa T, Kleckner N, Bishop DK (1997a) Saccharomyces cerevisiae recA homologues RAD51 and DMC1 have both distinct and overlapping roles in meiotic recombination Genes Cells 2:615–629

Shinohara A, Ogawa H, Ogawa T (1992) Rad51 protein involved in repair and recombination in S. cerevisiae is a RecA-like protein Cell 69:457–470

Shinohara A, Ogawa T (1998) Stimulation by Rad52 of yeast Rad51-mediated recombination Nature 391:404–407

Shinohara M, Gasior SL, Bishop DK, Shinohara A (2000) Tid1/Rdh54 promotes colocalization of Rad51 and Dmc1 during meiotic recombination Proc Natl Acad Sci U S A 97:10814–10819

Shinohara M, Oh SD, Hunter N, Shinohara A (2008) Crossover assurance and crossover interference are distinctly regulated by the ZMM proteins during yeast meiosis Nat Genet 40:299–309 doi:10.1038/ng.83

Shinohara M, Sakai K, Ogawa T, Shinohara A (2003) The mitotic DNA damage checkpoint proteins Rad17 and Rad24 are required for repair of double-strand breaks during meiosis in yeast Genetics 164:855–865

Shinohara M, Shita-Yamaguchi E, Buerstedde JM, Shinagawa H, Ogawa H, Shinohara A (1997b) Characterization of the roles of the Saccharomyces cerevisiae RAD54 gene and a homologue of RAD54, RDH54/TID1, in mitosis and meiosis Genetics 147:1545–1556

Smith AV, Roeder GS (1997) The yeast Red1 protein localizes to the cores of meiotic chromosomes J Cell Biol 136:957–967

Storlazzi A, Xu L, Cao L, Kleckner N (1995) Crossover and noncrossover recombination during meiosis: timing and pathway relationships Proc Natl Acad Sci U S A 92:8512–8516

Subramanian VV et al. (2016) Chromosome Synapsis Alleviates Mek1-Dependent Suppression of Meiotic DNA Repair PLoS biology 14:e1002369 doi:10.1371/journal.pbio.1002369

Sung P (1994) Catalysis of ATP-dependent homologous DNA pairing and strand exchange by yeast RAD51 protein Science 265:1241–1243

Sung P (1997) Function of yeast Rad52 protein as a mediator between replication protein A and the Rad51 recombinase The Journal of biological chemistry 272:28194–28197

Sym M, Engebrecht JA, Roeder GS (1993) ZIP1 is a synaptonemal complex protein required for meiotic chromosome synapsis Cell 72:365–378

Tang S, Wu MKY, Zhang R, Hunter N (2015) Pervasive and essential roles of the Top3-Rmi1 decatenase orchestrate recombination and facilitate chromosome segregation in meiosis Mol Cell 57:607–621 doi:10.1016/j.molcel.2015.01.021

Tsubouchi H, Argunhan B, Tsubouchi T (2018) Exiting prophase I: no clear boundary Current genetics 64:423–427 doi:10.1007/s00294-017-0771-y

Tsubouchi H, Roeder GS (2004) The budding yeast Mei5 and Sae3 proteins act together with Dmc1 during meiotic recombination Genetics 168:1219–1230 doi:10.1534/genetics.103.025700

Tsubouchi H, Roeder GS (2006) Budding yeast Hed1 down-regulates the mitotic recombination machinery when meiotic recombination is impaired Genes Dev 20:1766–1775 doi:10.1101/gad.1422506

Veaute X, Jeusset J, Soustelle C, Kowalczykowski SC, Le Cam E, Fabre F (2003) The Srs2 helicase prevents recombination by disrupting Rad51 nucleoprotein filaments Nature 423:309–312

Wu L, Hickson ID (2003) The Bloom’s syndrome helicase suppresses crossing over during homologous recombination Nature 426:870–874 doi:10.1038/nature02253

Xu L, Ajimura M, Padmore R, Klein C, Kleckner N (1995) NDT80, a meiosis-specific gene required for exit from pachytene in Saccharomyces cerevisiae Mol Cell Biol 15:6572–6581

Zakharyevich K, Ma Y, Tang S, Hwang PY, Boiteux S, Hunter N (2010) Temporally and biochemically distinct activities of Exo1 during meiosis: double-strand break resection and resolution of double Holliday junctions Mol Cell 40:1001–1015 doi:10.1016/j.molcel.2010.11.032

