## Supplemental Fig 1-5 and Table 1 for "Srs2 helicase prevents the formation of toxic DNA damage during late prophase I of yeast meiosis"

**Table S1: Strain list**

| Strain Name | Genotype |
|---|---|
| MSY832 | *MAT α, ho::LYS2, ura3, leu2::hisG, trp1::hisG, lys2* |
| MSY833 | *MAT a, ho::LYS2, ura3, leu2::hisG, trp1::hisG, lys2* |
| NKY1543 | *MAT α, ho::LYS2, ura3, leu2::hisG, lys2, his4X::LEU2-URA3, arg4-bgl* |
| NKY1303 | *MAT a, ho::LYS2, ura3, leu2::hisG, lys2, his4B::LEU2, arg4-nsp* |
| HSY315 | NKY1543, *srs2::TRP1, trp1::hisG* |
| HSY310 | NKY1303, *srs2::TRP1, trp1::hisG* |
| LPY071 | MYS832, *srs2::TRP1* |
| LPY072 | MSY833, *srs2::TRP1* |
| HSY596 | MSY 833 with *ndt80::LEU2* |
| HSY597 | MSY 832 with *ndt80::LEU2* |
| LPY058 | MSY 833 with *ndt80::LEU2*, *srs2::TRP1* |
| LPY059 | MSY 832 with *ndt80::LEU2*, *srs2::TRP1* |
| HSY185 | MSY 833 with *spo11-Y135F::KanMX6* |
| HSY186 | MSY 832 with *spo11-Y135F::KanMX6* |
| HSY462 | MSY 833 with *spo11-Y135F::KanMX6*, *srs2::TRP1* |
| HSY463 | MSY 832 with *spo11-Y135F::KanMX6*, *srs2::TRP1* |
| YFY74 | MSY 833 with *cdc20::pCLB2-CDC20::KanMX6* |
| YFY77 | MSY 832 with *cdc20::pCLB2-CDC20::KanMX6* |
| YFY80 | MSY 833 with *cdc20::pCLB2-CDC20::KanMX6*, *srs2::TRP1* |
| YFY83 | MSY 832 with *cdc20::pCLB2-CDC20::KanMX6*, *srs2::TRP1* |
| YFY03 | MSY 833 with *cdc20::pCLB2-SGS1::KanMX6* |
| YFY05 | MSY 832 with *cdc20::pCLB2- SGS1::KanMX6* |
| H7790 | *MAT a, ho:: LYS2, URA3, leu2::hisG, his3::hisG, trp1::hisG, lys2, fpr::KanMX4,*<br>*RPL13A-2xFKBP12::TRP1, tor1-1::HIS3, RAD54-FRB::KanMX6* |
| H7791 | *MAT α, ho:: LYS2, ura3, LEU2, his3::hisG, trp1::hisG, lys2, fpr::KanMX4*<br>*RPL13A-2xFKBP12::TRP1, tor1-1::HIS3, RAD54-FRB::KanMX6* |

| | |
|---|---|
| HYS82 | H7790 with *srs2::TRP1* |
| HYS71 | H7791 with *srs2::TRP1* |

**Supplemental Figure Legends**

**Supplemental Figure S1. Physical analysis of meiotic recombination in *srs2* mutant.**

A. Schematic representation of the *HIS4-LEU2* recombination hotspot.

B. DSB formation (top) and CON/NCO formation (bottom) at the *HIS4-LEU2* locus in the wild type and *srs2* strains were verified by Southern blotting. Genomic DNA was digested with *Pst*I for DSB and with XhoI+MluI for CON/NCO.

C. E. Kinetic analyses of meiotic DSBs and CO/NCO formation. Parental and DSB bands were quantified and % of DSB (top graph) and CO (second graph) or NCO (third graph) was calculated. Graphs show the mean values with variation (n=2). Wild type cells (blue circles; NKY1303/1543) and *srs2 cells* (red circles; HSY310/315).

D. F. Immunostaining analysis of a SC protein, Zip1 (green), was carried out in wild type and mutant cells. Representative images are shown for each strain. Wild type, MSY832/833; and *srs2* (HSY310/315). The bar indicates 2 μm.

E. G. Kinetics of SC formation. Zip1 staining in wild type and mutant cells was classified shown in (H). Zip1 staining in the wild type and mutant cells was classified as follows: dot (Class I, blue); partial linear (Class II, green); full SC (Class III, red). Spreads containing Zip1 lines were classified into two classes with less than 5 (class II, zygotene) and more than 5 (class III, pachytene) Zip1 dots. A minimum of 100 cells were analysed per time point. Wild type cells, MSY832/833*;* *srs2* cells, HSY310/315.

F. H. Kinetics of Zip1-polycomplexes in wild-type and *srs2* cells. The spreads containing Zip1-polycomplexes (arrow in F) were counted at each time point.

**Supplemental Figure S2. Rad52 and Hed1 staining in *srs2* mutant.**

A. Immunostaining analysis of Rad51 (green) and Dmc1 (red) on chromosome spreads in wild type (NKY1303/1543) and *srs2* (HSY310/315) mutant cells. Representative image with or without DAPI (blue) dye at 4 and 8 h for wild type and the *srs2* cells is shown. The bar indicates 2 μm.

B. Western blotting analysis of Dmc1 and Mei5 proteins during meiosis. Cell lysates at different time points in meiosis in wild type (NKY1303/1543) and *srs2* (HSY310/315) cells were probed with anti-Dmc1, anti-Mei5 and anti-tubulin.

C. Immunostaining analysis of Rad51 (green) and Rad52 (red) on chromosome spreads in wild type (NKY1303/1543) and *srs2* (HSY310/315) cells. Representative image with or without DAPI (blue) dye at 4 and 7 h for wild type and the *srs2* is shown. The bar indicates 2 μm.

D. Kinetics of Rad52 focus-positive cells in various yeast strains. The focus and aggregates were counted as shown in (D). Graphs show kinetics of one representative experiment for the wild-type cells (top; NKY1303/1543), and *srs2* cells (bottom; HSY310/315). Closed circles, Rad51 foci in wild type; open circles, Rad51 foci in *srs2*; open diamonds, Rad51 aggregates in *srs2*; closed triangles, Rad52 foci in wild type; open triangles, Rad52 foci in *srs2*; open square, Rad52 aggregates in *srs2*.

E. Immunostaining analysis of Hed1 (red) and Rad51 (green) on chromosome spreads in wild type (NKY1303/1543) and *srs2* (HSY310/315) cells. Representative image with or without DAPI (blue) dye at 4 and 8 h for wild type and the *srs2* is shown. The bar indicates 2 μm.

F. Kinetics of Rad51 or Hed1 focus-positive cells in various yeast strains. A spread with the foci of Rad51 or Hed1 is defined as a cell with more than five foci. Spreads containing Rad51/Hed1 aggregates were also counted. A minimum of 100 cells were analysed at each time point. Graphs show kinetics of one representative experiment for the wild type cells (top; NKY1303/1543), and *srs2* cells (bottom; HSY310/315). Closed circles, Rad51 foci in wild type; open circles, Rad51 foci in *srs2*; open diamonds, Rad51 aggregates in *srs2*; closed triangles, Hed1 foci in wild type; open triangles, Hed1 foci in *srs2*; open square, Hed1 aggregates in *srs2*.

**Supplemental Figure S3. Rad51 staining in *srs2 ndt80* mutant.**

A. Immunostaining analysis of Rad51 (red) and Nop1 (green) on chromosome spreads in *srs2* (HSY310/315) cells. Representative image with or without DAPI (blue) dye at 7 h for wild type and the *srs2* is shown. The bar indicates 2 μm.

B. Immuno-staining analysis of Rad51 in *spo11-Y135F srs2 (*HSY452/453) cells at 4 h.
C. Immunostaining analysis of Rad51 (green) and Dmc1 (red) on chromosome spreads in wild type (NKY1303/1543) and *SGS1-mn* (YFY03/05) cells. Representative images with or without DAPI (blue) dye at 4 and 7 h for wild type and the *srs2* cells are shown.
D. Kinetics of Rad51 foci-positive cells in *SGS1-mn* (YFY03/05) cells. The kinetics of Rad51 focus positive spreads were analyzed as shown in Figure 2B. Wild-type (closed circles); *SGS1-mn* (open circles).
E. Immunostaining analysis of Rad51 (green) on chromosome spreads in *ndt80* (HSY596/597) and *srs2 ndt80* (LPY058/059) cells. Representative image with or without DAPI (blue) dye at 4, 6, and 8 h for each strain is shown.
F. Kinetics of Rad51 focus-positive cells in *ndt80* (HSY596/597) and *srs2 ndt80* (LPY058/059) cells. Spreads containing Rad51 foci were counted also. A minimum of 100 cells were analysed at each time point. Graphs show kinetics of one independent experiment.

**Supplemental Figure S4. Immuno-staining analysis of Rad51 and Zip1/Rec8 in *srs2 CDC20mn* cells.**

A. Immuno-staining analysis of Rad51 and Zip11 in wild type (NKY1303/1543) and *srs2* (HSY310/315) cells. The chromosome spreads immuno-stained against Rad51 (green) as well as chromosome protein Zip1 (red) are shown.
B. Immuno-staining analysis of Rad51 and Red1 in wild type (NKY1303/1543) and *srs2* (HSY310/315) cells. The chromosome spreads immuno-stained against Rad51 (green) as well as chromosome protein Red1 (red) are shown.
C. Kinetics of Rad51 aggregate-positive cells in Red1-postive spreads. Rad51-focus and Rad51-aggregate positive spreads were classified into Red1-negative (open bars) and Red1-positive (closed bars) at each time point.
D. Immuno-staining analysis of Rad51 and Rec8 in wild type (NKY1303/1543) and *srs2* (HSY310/315) cells. The chromosome spreads immuno-stained against Rad51 (green) as well as chromosome protein Rec8 (red) are shown.

E. Immuno-staining analysis of Rad51 and Rec8 in *CDC20mn* (YFY74/77) and *srs2 CDC20mn* (YFY80/83) cells. The chromosome spreads immuno-stained against Rad51 (green) as well as chromosome protein Rec8 (red) are shown.

**Supplemental Figure 5. CHEF and CHEF-Southern blotting**

EtBr-staining of CHEF analysis of yeast chromosomes during meiosis. Chromosomal DNAs in yeast cells of wild type (NKY1303/1543) and *srs2* (HSY310/315) at each time point of meiosis were analyzed by CHEF gel electrophoresis and after the electrophoresis, the gels were stained with EtBr.

### Figure S1. Sasanuma/Sakurai

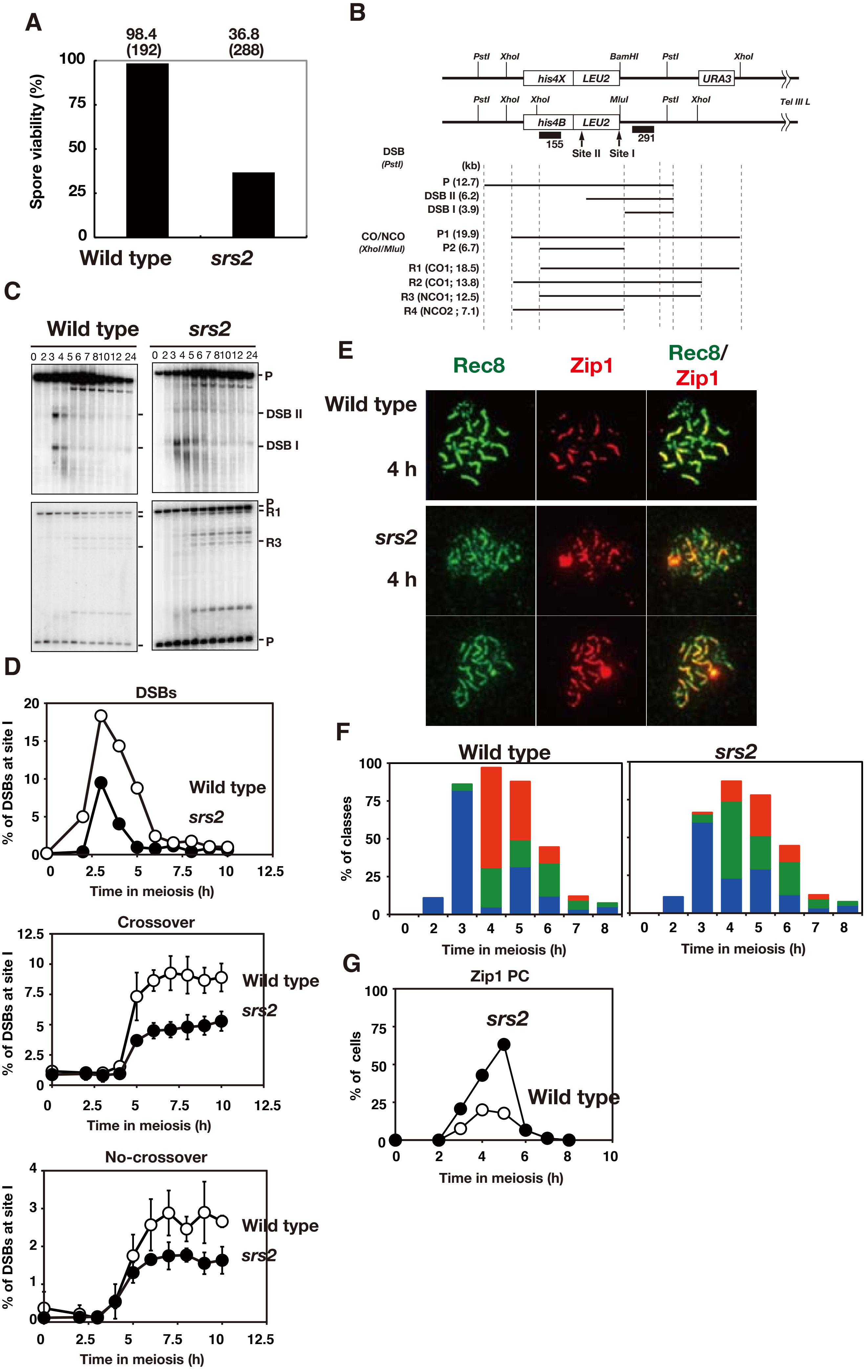

Figure S2. Sasanuma/Sakurai

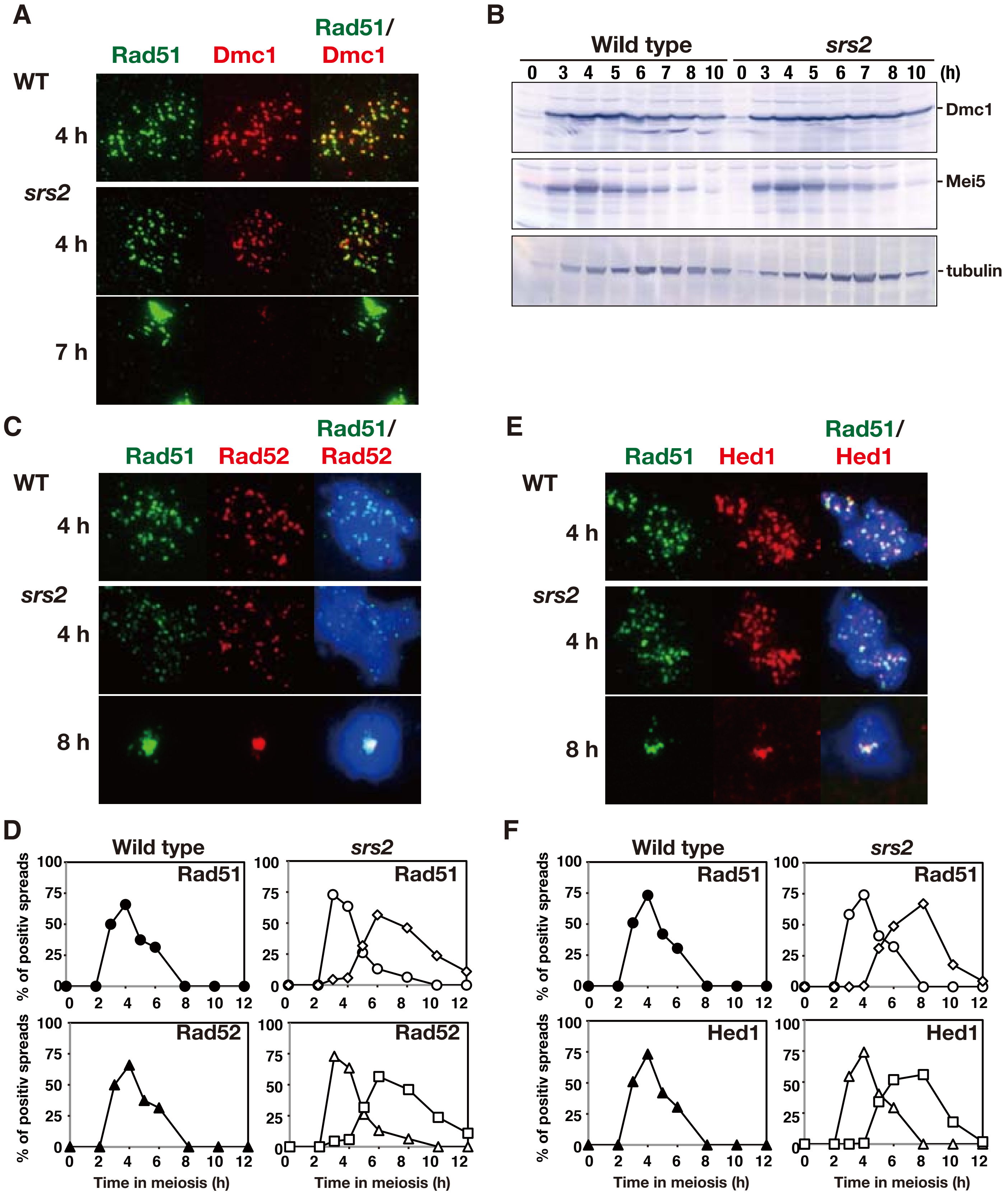

### Figure S3. Sasanuma/Sakurai

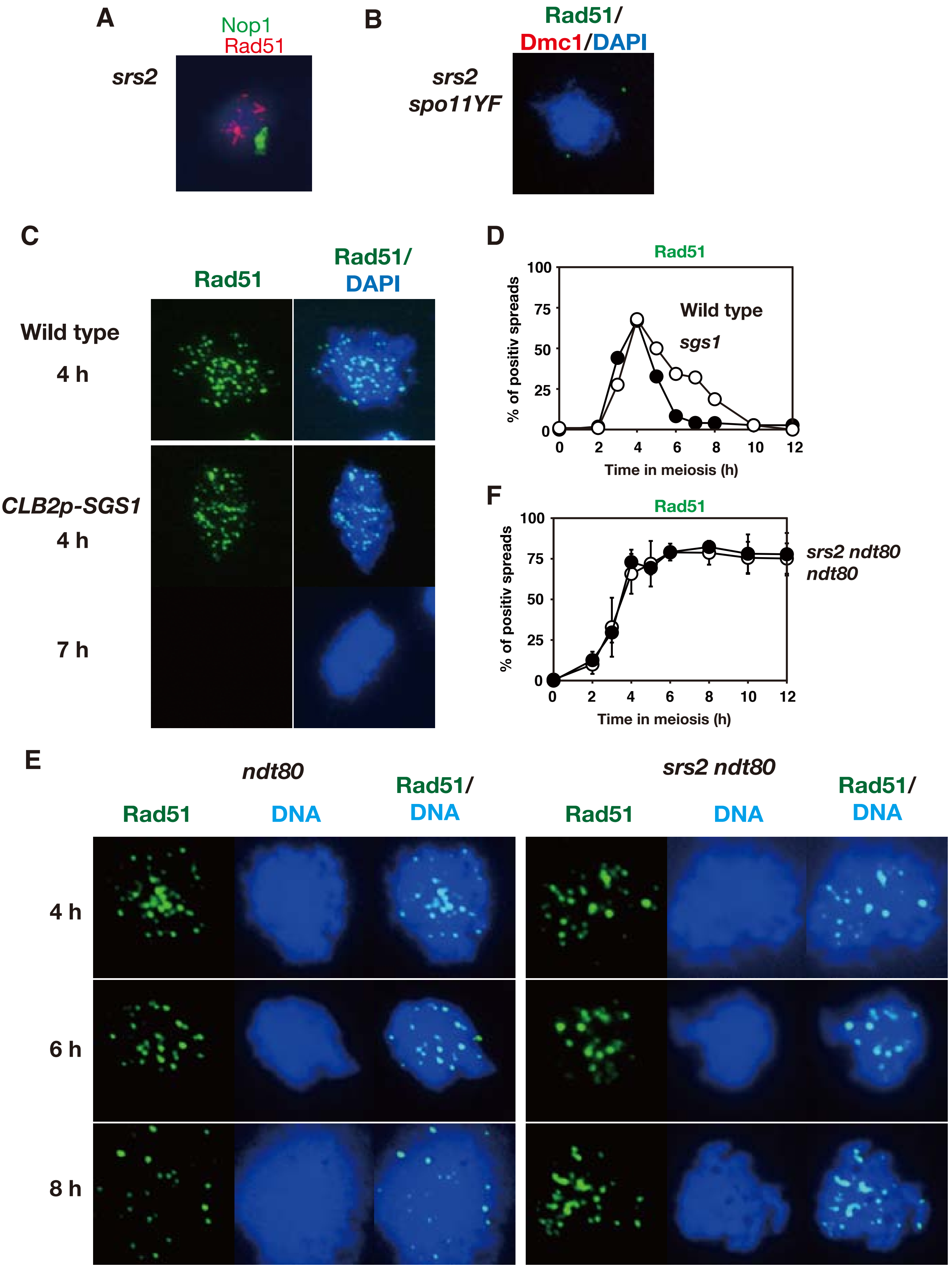

Figure S4. Sasanuma/Sakurai

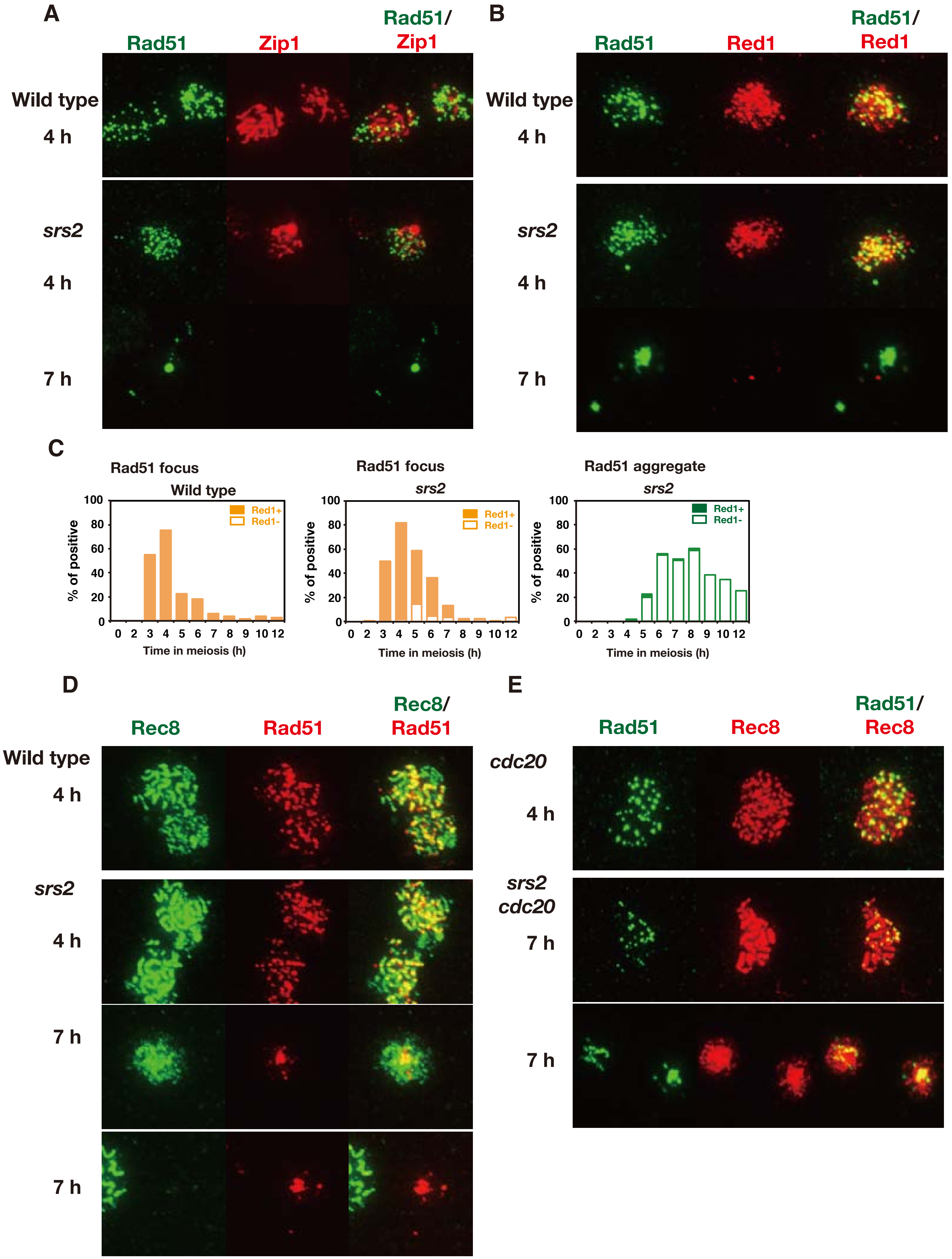

### Figure S5. Sasanuma/Sakurai

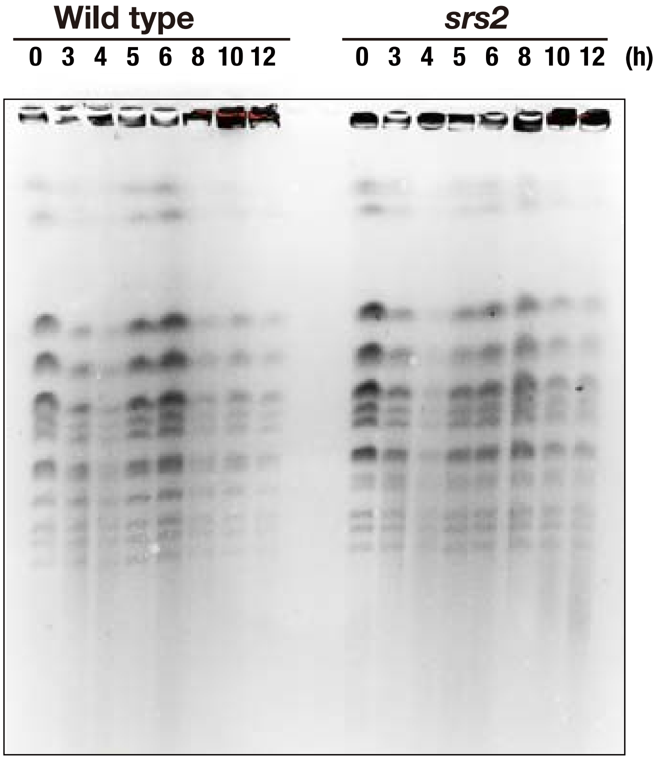
